# Spatial modulation of hippocampal activity in freely moving macaques

**DOI:** 10.1101/2020.10.03.324848

**Authors:** D. Mao, E. Avila, B. Caziot, J. Laurens, J.D. Dickman, D.E. Angelaki

## Abstract

The hippocampal formation is linked to spatial navigation, but there is little corroboration from freely-moving primates with concurrent monitoring of three-dimensional head and gaze stances. We recorded neurons and local field potentials across hippocampal regions in rhesus macaques during free foraging in an open environment while tracking their head and eye. Theta band activity was intermittently present at movement onset and modulated by saccades. Many cells were phase-locked to theta, with few showing theta phase precession. Most hippocampal neurons encoded a mixture of spatial variables beyond place fields and a negligible number showed prominent grid tuning. Spatial representations were dominated by facing location and allocentric direction, mostly in head, rather than gaze, coordinates. Importantly, eye movements strongly modulated neural activity in all regions. These findings reveal that the macaque hippocampal formation represents three-dimensional space using a multiplexed code, with head orientation and eye movement properties dominating over simple place and grid coding during free exploration.

## Introduction

The rodent hippocampal formation (HF) is implicated in spatial navigation. At the network level, hippocampal local field potential (LFP) shows prominent theta oscillations (6-10 Hz) during movement (Vanderwolf, 1969). Theta power and frequency are correlated with locomotion speed (McFarland et al., 1975; Sławińska and Kasicki, 1998), suggestive of a role of theta oscillations in spatial navigation. The precise relationship between theta phase and neuronal firing reflects a temporal code that may support communication across regions and organize ensemble activity into sequences (Mizuseki et al., 2009; O’Keefe and Recce, 1993; Skaggs et al., 1996). In contrast to rodents, primate studies have consistently found a lack of continuous theta oscillations either during virtual navigation or real world locomotion (Ekstrom et al., 2005; Talakoub et al., 2019), but spike-LFP phase coding appears to be conserved to some extent, at least in humans (Jacobs et al., 2007; Qasim et al., 2020).

At the cellular level, multiple cell types have been identified to encode position, head direction, speed and other spatial variables in the rodent HF (Hafting et al., 2005; McNaughton et al., 1983; O’Keefe and Dostrovsky, 1971; Taube et al., 1990), some with 3D properties (Angelaki et al., 2020; Grieves et al., 2020), that has also been found in the bat HF (Finkelstein et al., 2015; Yartsev and Ulanovsky, 2013). Recent studies have employed multimodal models to reveal multiplexed representations in the rodent HF (Hardcastle et al., 2017; Laurens et al., 2019; Ledergerber et al., 2020). A fundamental advantage of such models is that they can correctly identify multimodal responses even for variables that are correlated and interdependent, while proved remarkably immune to pitfalls such as overfitting; whereas traditional methods often fail to quantify neurons with mixed selectivity (Laurens et al., 2019).

The primate HF has been scarcely explored under the same, freely-moving paradigms. Human studies have been mostly limited to virtual navigation in patients and focused exclusively on a limited number of spatial variables, including virtual position and direction (Doeller et al., 2010; Ekstrom et al., 2003; Jacobs et al., 2013; Maguire et al., 1998). A recent study also found LFP correlates of proximity to environmental boundaries in mobile humans (Stangl et al., 2020). Non-human primate studies have used almost exclusively head-fixed monkeys, either during cart navigation (Matsumura et al., 1999; O’Mara et al., 1994) or in a virtual environment setting (Furuya et al., 2014; Gulli et al., 2020; Wirth et al., 2017). In these studies, hippocampal neurons showed correlates with self-location, attended location, and head direction. Three limited investigations in freely-moving monkeys have reported location-specific hippocampal activity reminiscent of rodent place cells albeit more dispersed and less prevalent (Courellis et al., 2019; Hazama and Tamura, 2019; Ludvig et al., 2004). Yet, these studies neither explored three-dimensional properties nor used contemporary multimodal models to quantify multi-variable spatial representations.

The primate HF also shows neural correlates with visual space (Nau et al., 2018; Rolls, 2021). A rather provocative observation is the existence of putative gaze-centered spatial representations in the HF of head-fixed macaques (Killian et al., 2012; Meister and Buffalo, 2018; Wirth et al., 2017), which may imply differences between primates and rodents. Of particular relevance is the report that some hippocampal neurons showed correlates with the location that the animal was looking at (‘spatial view’ cells) but not where the animal was located (Georges-François et al., 1999; Rolls et al., 1997). This property, however, has yet to be interrogated by monitoring both eye and head movements in truly freely-moving primates where flexible head movement is also a critical component of natural behavior.

Here, for the first time, we combine precise head tracking, wireless eye tracking, and telemetric recordings across three HF regions – hippocampus, entorhinal cortex, and subicular complex, in freely-foraging macaques as they explore a three-dimensional environment.

## Results

To investigate how the primate HF represents space during ambulatory navigation, we trained three macaques to freely forage in an open circular arena endowed with salient cues (Fig. 1A). Using chronically implanted tetrodes or single electrodes (Fig. 1B,C, and S1), we recorded LFP and single neuron activity across three HF regions in both hemispheres: hippocampus (HPC), entorhinal cortex (EC), and subicular complex (SUB) (HPC, EC and SUB in monkeys K and L, only EC in monkey B). To reconstruct electrode locations, we co-registered preoperative magnetic resonance imaging (MRI) images and postoperative computed tomography (CT) images (Fig. 1D-G and S1), corroborated by histology in one monkey (Fig. S1 bottom right). To accurately track the monkeys’ head movement, we used a marker-based approach, combined with wireless eye tracking in one animal (Fig. 1H,I). Many spatial variables were extracted from the markers’ trajectories in 3D space (Fig. 1J and S2). Overall, the monkeys’ behavior covered a broad space while salient landmarks had a strong influence (Fig. S3). Wireless eye tracking revealed eye-in-head movements were predominantly within ±30° horizontal and ±20° vertical (Fig. S3A bottom right).

**Fig. 1.**
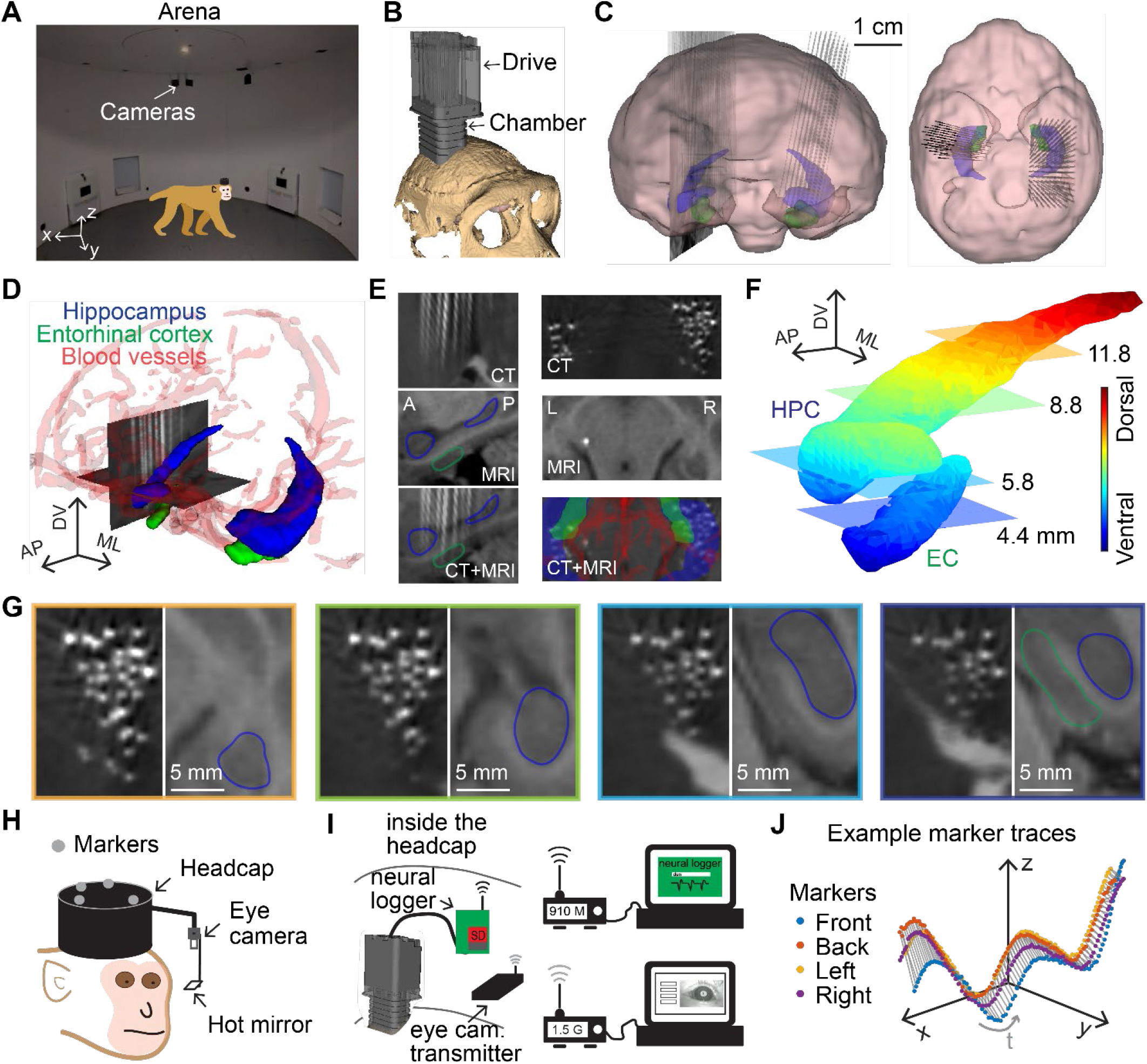
Freely-moving monkey setup and electrode localization. (A) Monkeys were trained to freely forage for randomly scattered food pellets or fruits in an open circular arena (pictured, 330 cm diameter, 212 cm height) equipped with a motion tracking system. (B) A large-scale semi-chronic microdrive holding 124 independently movable electrodes (42 mm travel distance) was implanted in the right hemisphere of monkey K. The drive and skull models are shown. (C) Segmented models of the brain, where hippocampus (blue) and entorhinal cortex (green) are shown (front and top views). (D) Segmented models of the hippocampus (blue), entorhinal cortex (green), and blood vessels (red). Co-registered CT images are shown. White tracks are electrodes. (E) CT, MRI, and co-registered CT and MRI images for sagittal (left) and axial (right) views. White tracks and dots are electrodes. (F) Models of the hippocampus and entorhinal cortex color-coded by dorsal-ventral position. Four axial planes are highlighted. (G) CT and MRI images corresponding to the 4 axial plans shown in (F). Hippocampus is circled in blue; entorhinal cortex is circled in green. Electrodes are shown as white dots. Deeper electrodes are also visible in more dorsal planes. (H) A marker-based approach was used to track head motion in 3D. Four markers were placed on the head cap. An eye tracker was attached to the head ring for tracking the pupil position of the left eye of monkey K. (I) Neural data were recorded and stored in a neural logger that was regularly synchronized with the computer. Eye tracking images were wirelessly transmitted. (J) A short segment of the markers’ trajectories in 3D, from which head position, translation, and orientation were extracted. See also **Fig. S1-S3**.

### Speed-dependent low theta activity is linked to movement onset

We first examined the relationship between hippocampal LFP and behavior. Hippocampus showed stronger absolute power than the EC and SUB (Fig. 2A) (power comparison for 1-10 Hz and 10-50 Hz, both p < 0.001, one-way ANOVA). After removing aperiodic activity (Donoghue et al., 2020), the LFP showed two prominent oscillatory components: one peaked in the low theta band (1-4 Hz) and another in the beta band (12-30 Hz) (Fig. 2A). The HPC showed the strongest oscillatory power in the low theta band but the weakest in the beta band (Fig. 2A) (both p < 0.001, one-way ANOVA). In sharp contrast to rodents, and consistent with another freely-moving monkey study (Talakoub et al., 2019), hippocampal LFP lacked a sustained oscillatory pattern in the theta band during locomotion (Fig. 2B).

**Fig. 2.**
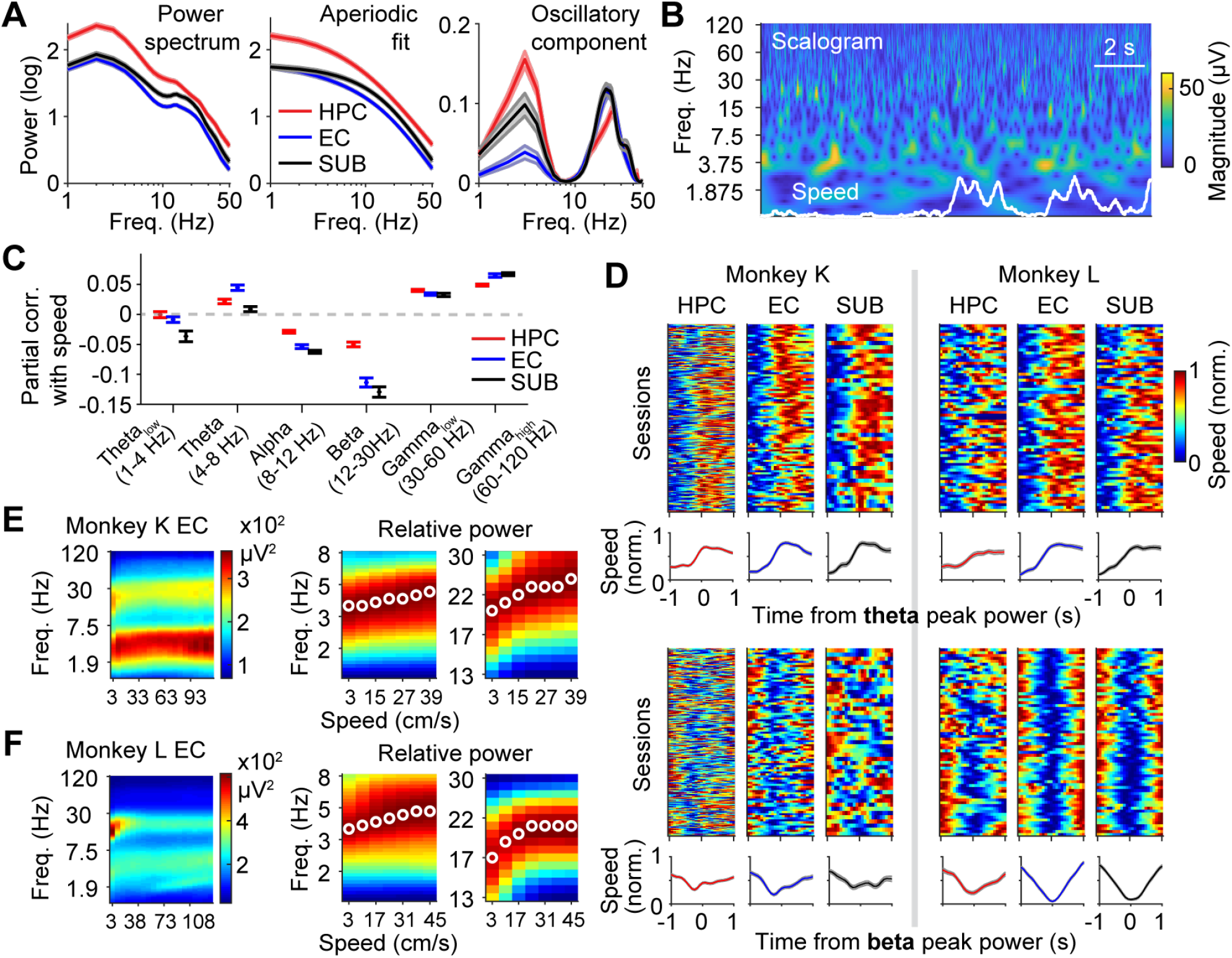
Speed modulation of hippocampal local field potential. (A) Left, absolute power spectrum; middle, aperiodic fit to the left; right, power spectrum of the oscillatory component (absolute - aperiodic). Shaded areas: SEM over sessions. Data are average across sessions/animals K and L, unless otherwise indicated. (B) Scalogram of a short segment of hippocampal LFP. Corresponding speed trace is shown. (C) Mean partial correlation between LFP power in different bands and speed. Error bars: SEM over sessions. (D) Colormaps of peak theta (top) and beta (bottom) power-triggered, normalized speed, shown for each session separately and as mean±SEM across sessions. (E) Left, colormap of LFP power plotted in frequency vs. speed for monkey K right EC. Middle, relative power (normalized such that the power ranges from 0 to 1 in each speed bin) in the theta and beta bands. White dots indicate the peak frequency in each speed bin. (F) The same as (E) for monkey L left EC. These effects were not as obvious in other regions.

To assess the correlation between locomotion speed and LFP power in different bands, we employed a wavelet-based approach, which is ideal to capture localized variations of power. We calculated the partial correlations between LFP power in 6 bands and locomotion speed, by taking into consideration other behavioral variables (Fig. 2B and S2). Theta (4-8 Hz), low (30-60 Hz) and high (60-120 Hz) gamma power showed an overall positive correlation with speed, whereas alpha (8-12 Hz) and beta (12-30 Hz) activity showed a negative correlation across HF regions (Fig. 2C).

Intermittent theta activity has been reported, but it is unclear what the intermittent theta activity correlates with during free behavior. By examining theta-triggered speed profiles, we found that theta power was linked to movement onset across the HPC, EC, and SUB, in sharp contrast to the opposite effect of speed on beta power (Fig. 2D). Both theta and beta frequencies increased with speed at slow motion, most obvious for the EC (Fig. 2E,F).

### Hippocampal neurons are tuned to multiple spatial variables

Both recording methods, single electrodes and tetrodes, allowed good single unit isolation (Fig. 3A,B). We focused on the 599 well-isolated single units: HPC, 273 neurons; EC, 216 neurons; SUB, 110 neurons (monkey K: HPC 186, EC 49, SUB 27, n = 37 sessions; monkey L: HPC 87, EC 137, SUB 83, n = 86 sessions; monkey B: EC 30, n = 13 sessions). Spike amplitude ranged from 64 to 124 μV (20-80 percentiles) (Fig. 3C). Recordings were stable throughout the sessions (spike feature correlation between the 1^st^ half and the 2^nd^ half of sessions: Pearson’s r = 0.997, p < 0.001) (Fig. 3D). We used independent measures to verify spike sorting quality (Fig. 3E,F).

**Fig. 3.**
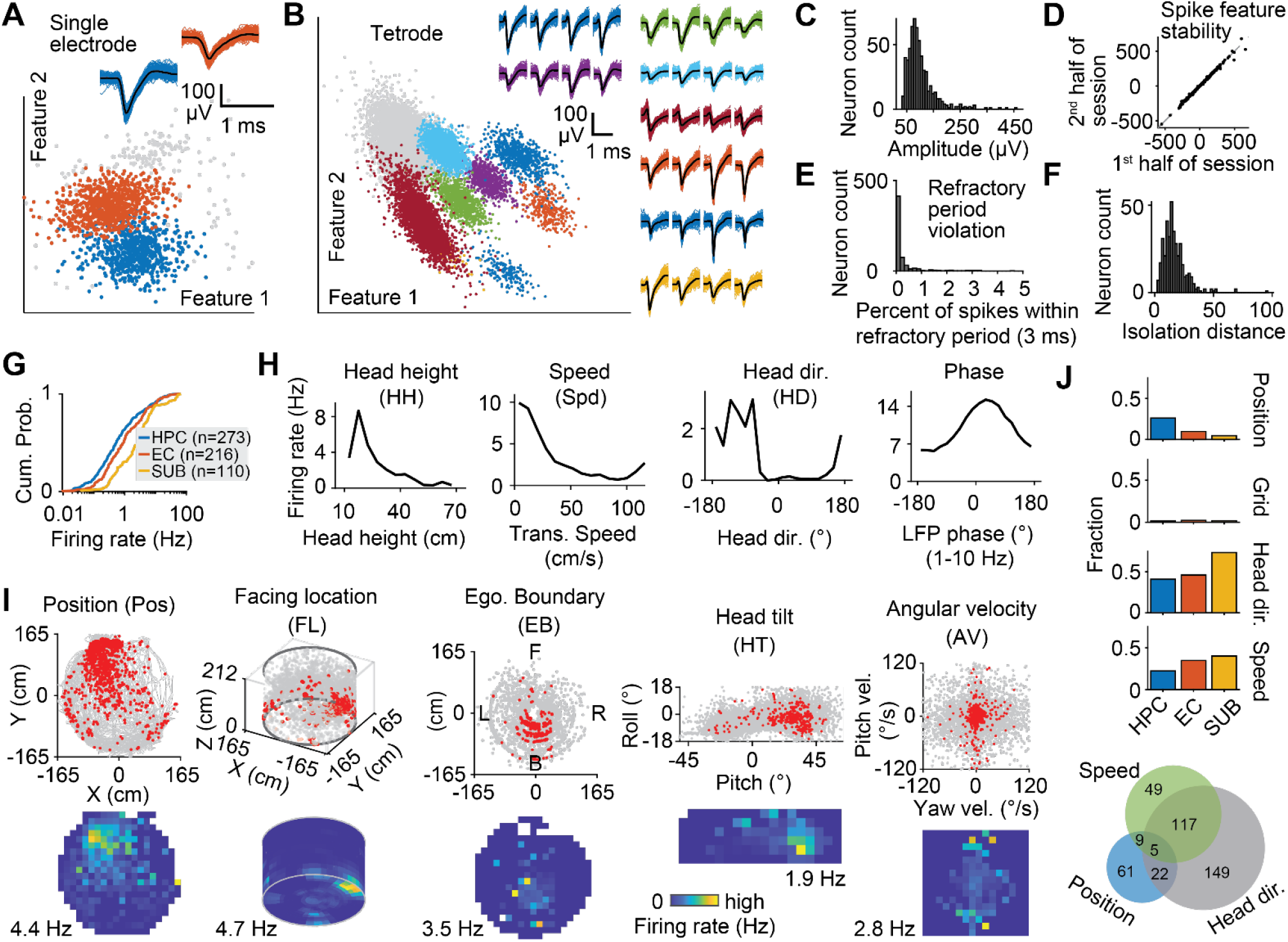
Diverse spatial tuning across hippocampal regions. (A) Spike sorting for an example single electrode (2 isolated units). (B) Spike sorting for an example tetrode (8 units). (C) Spike amplitude distribution for all single units. (D) Spike feature correlation between the 1^st^ half and the 2^nd^ half of all sessions. (E) Distribution of percent of spikes within refractory period (3 ms) for all single units, i.e. refractory period violation; For 72% of the neurons the violation was less than 0.2%. (F) Distribution of isolation distance for all qualified single units (Methods). (G) Cumulative distribution of the mean firing rates for neurons recorded across three HF regions. (H) Example raw tuning curves plotted as the average firing rate vs. head height, translational speed, azimuth head direction, and LFP phase (1-10 Hz). (I) Example raw data (top) and raw tuning curves (bottom) for horizontal position (place fields), facing location (where the head points on the arena surface), egocentric boundary, head tilt, and angular velocity. Red dots: spikes; gray: behavioral variables, down-sampled for visualization. Tuning curves (bottom) show colormaps of raw firing rate as a function of each spatial variable. In each panel, the peak firing rate (yellow) is indicated with a number at the bottom of the colormaps. The lowest firing rate (dark blue) is 0 for all panels. The variable abbreviations are indicated in the title for each panel in (H) and (I). Tuning curves are all from different neurons. (J) Fraction of neurons modulated by the most commonly used spatial variables (position, grid, azimuth head direction, and speed) for each region, computed using traditional analyses (Methods). Bottom, Venn diagram showing the number of neurons encoding each variable and conjunctive variables. See also **Fig. S2, S4 and S5**.

Consistent with previous studies (Barnes et al., 1990; Skaggs et al., 2007), the activity of HF neurons was in general sparse and the firing rates showed log-normal distributions with SUB neurons showing the least sparsity (firing rate: HPC, 0.14-3.05 Hz; EC, 0.24-4.41 Hz; SUB, 0.56-7.15 Hz; 20-80 percentiles; p < 0.001, Kruskal-Wallis test) (Fig. 3G). Individual HF neurons exhibited tuning to diverse spatial variables, including horizontal position, head height, linear speed, azimuth head direction, head tilt, head facing location (where the head points, a 3D variable), egocentric boundary (relative position and direction to arena boundary), and head angular velocity (Fig. 3H,I, S2 and S4).

We first used traditional analyses to quantify neuronal tuning for the four most extensively studied variables in rodents: position, grid, azimuth head direction and linear speed,. Based on such analyses, which compared data against permutated distributions (Fig. S5; Methods), hippocampal neurons showed prominent modulation by position (HPC, 26%; EC, 10%; SUB, 5%), speed (HPC, 22%; EC, 35%; SUB, 40%) and head direction (HPC, 41%; EC, 46%; SUB, 74%) (Fig. 3J). A large fraction of neurons (153/599, 26%) showed conjunctive tuning (Fig. 3J, bottom). The proportion of cells with significant grid modulation was rather low (at 99%: HPC 1% 3/273, EC 2% 4/216, SUB 2% 2/110; see Fig. S5C for an example). Dropping the significance criterion barely increased these numbers: at 95%: HPC 6% (16 cells), EC 6% (13 cells), SUB 3% (3 cells). Thus, other than the absence of notable grid tuning, macaque and rodent HF appear similar when traditional analysis is used.

### Mixed selectivity across hippocampal regions is dominated by head facing location

One issue with the traditional analyses is that a neuron may appear to encode a particular variable when other more relevant variables are excluded, therefore masking the actual, more multiplexed coding schemes. Several recent studies have emphasized the advantages of using multimodal models to characterize mixed selectivity, which are agnostic to tuning curve shape and robust to the interdependence of encoded variables (Hardcastle et al., 2017; Laurens et al., 2019).

To quantify the simultaneous encoding of many spatial variables by HF neurons in an unbiased way, we used a variant of the cross-validated, multivariate linear-nonlinear Poisson (LNP) model framework (Fig. 4A; Methods) (Laurens et al., 2019). A variable was encoded if its inclusion significantly improved the model’s performance (p < 0.05, one-sided Wilcoxon signed rank test; Methods) (Fig. 4A and S6). Thus, each neuron could encode a single or combination of variables – or not be tuned to any spatial variable at all.

**Fig. 4.**
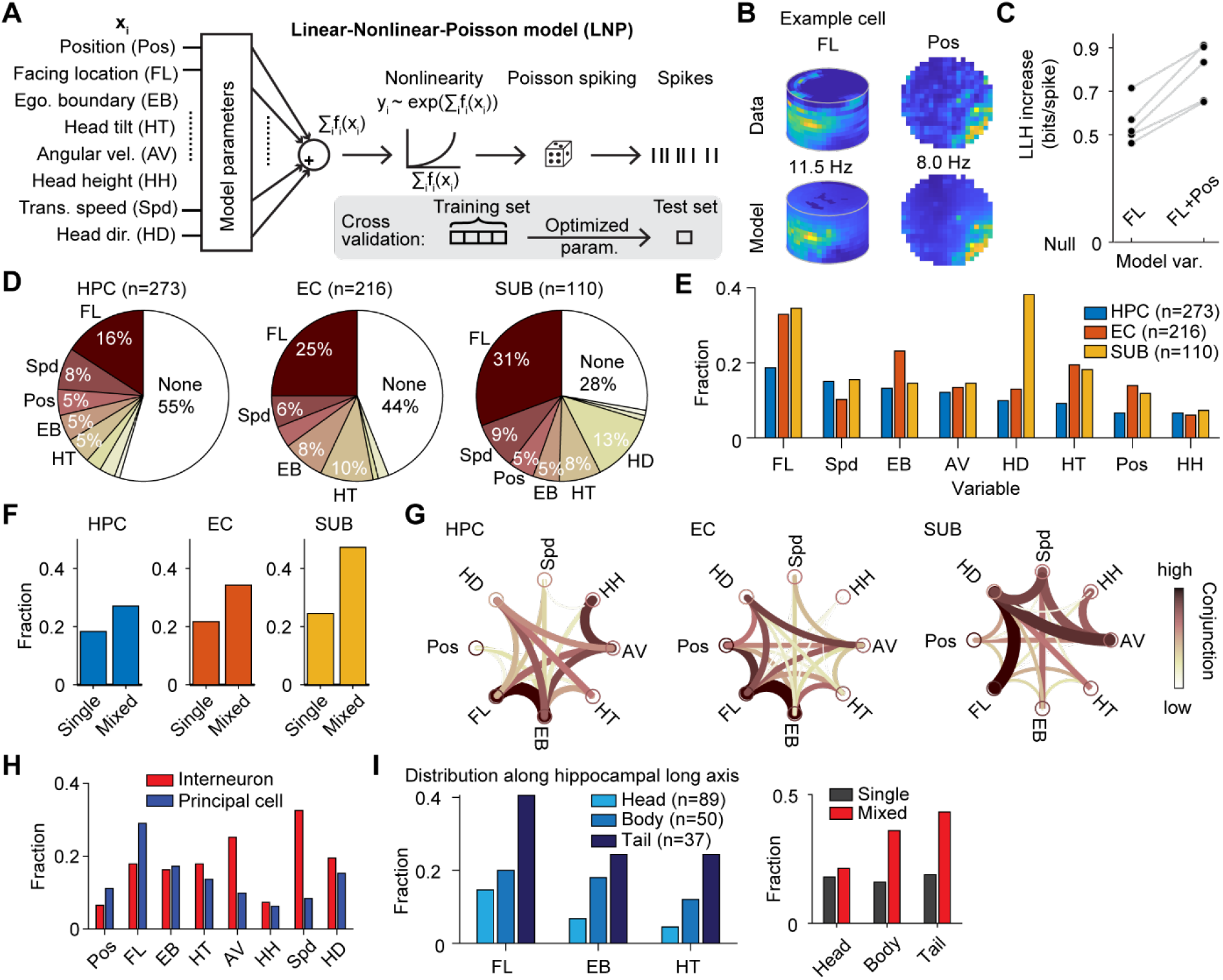
Mixed selectivity of diverse spatial variables. (A) A model-based statistical framework was used to quantify the spatial coding. Any single variable or combination of multiple variables was fitted to the data. A forward search approach was used to select the best model. A 5-fold cross-validation was used to test the significance of model fitting (Methods). (B) An example cell showing tuning to facing location and position. (C) Log-likelihood (LLH; goodness of fit) increase as a function of model variables for the example cell shown in (B). (D) All neurons were classified according to the single variable that best explained their responses. Pie chart shows the name and the fraction of select variables that were best encoded in each region. ‘None’ cluster corresponds to neurons that did not encode any of these variables (no spatial tuning model was better than the null model; Methods). (E) Breakdown of the fraction of encoding neurons for each variable in each region. A single neuron can encode none, one, or more than one variable. (F) Fraction of neurons tuned to single variable (single) or to more than one variable (mixed) in each region. (G) Circular graph representation of the degree of conjunction between variables. Line thickness and color correspond to how often two variables are co-coded in a single neuron. The thicker and the darker the line, the stronger the conjunction between the two variables. (H) Fraction of putative interneurons and principal cells encoding each spatial variable. (I) Left, fraction of encoding neurons along the hippocampal long axis for variables FL, EB, and HT. Right, fraction of neurons showing single and mixed encoding along the hippocampal long axis. See also **Fig. S4, S6, and S7**.

The main model included 8 variables: 3D position, which was decomposed into 2D position (Pos) and head height (HH), angular velocity (AV), linear speed (Spd), egocentric boundary (EB), facing location (FL), as well as 3D orientation, which was decomposed into head tilt (HT) and azimuth head direction (HD) (Fig. 4A and S2; Methods). Note that FL is different from 3D orientation: Depending on the animal’s position, the same head orientation can yield distinct FL, and different head orientations can point to the same FL. Overall, this model provided good fit to the data (Fig. 4B,C), with the SUB exhibiting the strongest encoding (fraction of neurons encoding at least one of these variables: HPC: 45%, EC: 56%, SUB: 72%; Fig. 4D). Overall, HF population exhibited rich coding schemes (Fig. 4 and S6).

We first classified neurons based on the single variable that best explained their firing. Across all HF regions, the most frequently coded variable was FL (22%). This variable is similar to ‘Spatial View’ previously reported to dominate the macaque HPC and parahippocampal tuning (Rolls et al., 1997). FL was more prevalent in the SUB (31%) and EC (25%) than HPC (16%) (Fig. 4D). Other dominant variables represented were linear speed (HPC: 8%, EC: 6%, SUB: 9%), EB (HPC: 5%, EC: 8%, SUB: 5%) and 3D head orientation, comprised of HD and HT. HD was a dominant coded variable mainly in the SUB (13%), whereas HT (vertical orientation tuning) was more broadly coded in the EC and SUB. In contrast to the dominance of orientation variables, only 6% of HPC cells were tuned to either component of 3D position (horizontal Pos: 5%; HH: 1%). The proportion of position-tuned neurons was also low for EC (4%; Pos: 4%; HH: 0%) and SUB (6%; Pos: 5%; HH: 1%). Note that Pos tuning encompasses not only place, but also grid and border fields.

Considering simultaneous encoding of multiple variables, FL was again the most prominently coded variable across HF regions (HPC: 19%, EC: 33%, SUB: 35%) (Fig. 4E and S7). Notably, this variable was coded in a higher percentage of EC/SUB than HPC cells (Fig. 4E). Mixed selectivity was ubiquitous across all HF regions (single vs. mixed selectivity, HPC: 18% vs 27%, EC: 22% vs 34%, SUB: 25% vs 47%; Fig. 4F), and it was not explained by spike sorting quality (Fig. S4C,D).

Overall we found that all spatial variables examined were represented in the macaque HF. When mixed selectivity was taken into account, the percent of cells tuned to linear speed doubled. A similar percentage of neurons was also tuned to angular velocity (Fig. 4E). Thus, angular velocity and linear speed were often co-coded together with other spatial variables. This conclusion also applies to EB, which was represented in ~20% of the population, most prominently encoded by EC neurons (23%; Fig. 4E). Notably, even after considering mixed selectivity, position coding was relatively weak, particularly in the HPC (7%; EC: 14%; SUB: 12%). Considering together Pos and HH, 3D position tuning was also relatively infrequent compared to FL (HPC: 13%, EC: 19%, SUB: 17%). As in rodents, HD was most prominently coded in the SUB (38%). Considered together with HT, a large proportion of cells was tuned to 3D head direction (HPC: 16%; EC: 28%; SUB: 48%).

Different HF regions carried different combinations of mixed selectivity (Fig. 4G). In both the HPC and EC, FL and EB were most often co-coded in the same neurons. In the EC, FL was also often coded with position and/or HD and AV. In contrast, speed tuning was often encountered in isolation in the HPC and EC, likely represented selectively in interneurons (see below). Finally, the dominant HD tuning was most often coded together with FL, AV and speed in the SUB (Fig. 4G). There was no geometric relationship in the preferred direction among FL, position, and HD tuning (Fig. S7C).

Spatial coding was present in both principal cells and interneurons, classified according to spike waveform (Barthó et al., 2004). We classified all neurons into narrow-spiking (NS, putative interneurons, n = 123, 21%) and broad-spiking (BS, putative principal cells, n = 476, 79%) cells (Fig. S4E,F). Linear speed and angular velocity tuning were more abundant in interneurons (Fig. 4H). In contrast, FL and Pos tuning were more common for principal cells.

Spatial coding in the rodent HF is topographically organized along the dorsal-ventral axis (Hafting et al., 2005; Jung et al., 1994). The monkey hippocampal rostral-caudal axis is analogous to the ventral-dorsal axis in rodents. In one monkey, we recorded from nearly the entire hippocampus in the right hemisphere (Fig. 1G and S1). We observed a gradual increase in the fraction of spatially-tuned neurons for the most prominent spatial variables: FL, EB, and HT (FL: p = 0.005; EB: p = 0.018; HT: p = 0.005, chi-square test) (Fig. 4I left). Mixed selectivity also increased from rostral to caudal hippocampus (p = 0.029, chi-square test) (Fig. 4I right).

These results show that, like rodents, a large proportion of HF neurons in the freely-moving macaques are tuned to spatial variables, and the majority show mixed selectivity. The two species are also similar in the topography of spatial coding along the hippocampal long axis. However, unlike the dominance of place coding in rodents, orientation-related variables dominate the macaque HF.

### Heterogeneous spatial coding covers the entire space

Spatial representations in the macaque HF are heterogeneous with distributed preferred firing fields, as shown for FL and HT tuning examples (Fig. 5A,B). Preferred fields for FL, HT, Pos and EB covered a broad space, although salient arena cues influenced the clustering at certain locations (Fig. 5C-F). Two major clusters were found for speed tuning: monotonically increasing or decreasing (Fig. 5G). Preferred HD uniformly tiled the azimuth directional space (p = 0.9, Rayleigh test), even though occupancy was strongly biased toward the direction of the entrance/exit door (Fig. 5H). HH tuning also spread across different head elevations, whereas most of AV-tuned neurons preferred firing at either low or high head angular velocities (Fig. S7D).

**Fig. 5.**
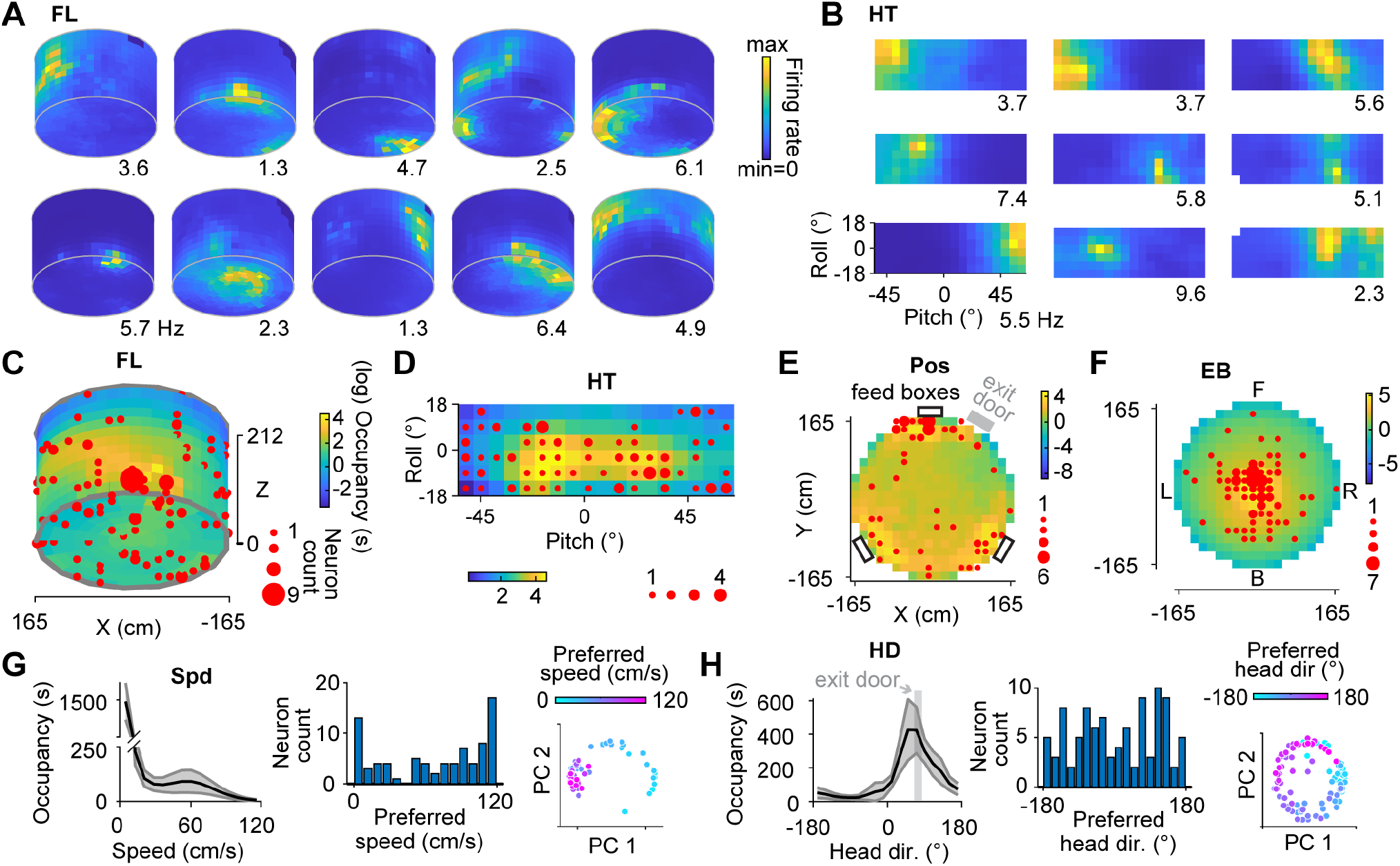
Spatial representations in the hippocampal formation are heterogeneous. (A) Example model-based tuning curves for 10 neurons coding facing location (FL). Peak firing rates are indicated. Darkest blue corresponds to bins that were not occupied. (B) Example model-based tuning curves for 9 neurons that encoded head tilt (HT). (C) Preferred firing locations (red dots) for all neurons encoding FL superimposed on the average occupancy colormap (log scale) across all monkeys. Dot size corresponds to the number of neurons. (D) Preferred firing fields (red dots) for all neurons encoding HT superimposed on the average occupancy colormap across all monkeys. Dot size corresponds to the number of neurons. (E) and (F) The same as (C) and (D) for position encoding neurons and egocentric boundary encoding neurons, respectively. Feed boxes are indicated as black rectangles. Exit/entrance door is indicated as gray rectangle in (E). (G) Left, average occupancy as a function of translational speed across monkeys. Shaded area indicates 1x standard deviation across sessions. Middle, distribution of preferred speed. Right, results of clustering analysis for the normalized speed tuning curves using principal component analysis (the first 2 components are shown). Each dot represents a single neuron. Colors correspond to their preferred speed. (H) The same as (G) for azimuth head direction. Gray bar indicates the direction of the exit/entrance door. See also **Fig. S7**.

### Spike-LFP phase-locking and phase precession

A large number of HF neurons were phase-locked to the LFP, and this was true across frequency bands (Fig. 6A,B) (low theta: 56%, theta: 25%, alpha: 25%, beta: 38%, low gamma: 58%, high gamma: 80%). Except for low theta, where preferred phases were more distributed, neurons preferred firing around LFP troughs (phase: ±180°) for all other frequency bands (Fig. 6B). Phase-locking was more common for spatially-tuned neurons (Fig. 6C), and no difference was seen for cells coding different variables.

**Fig. 6.**
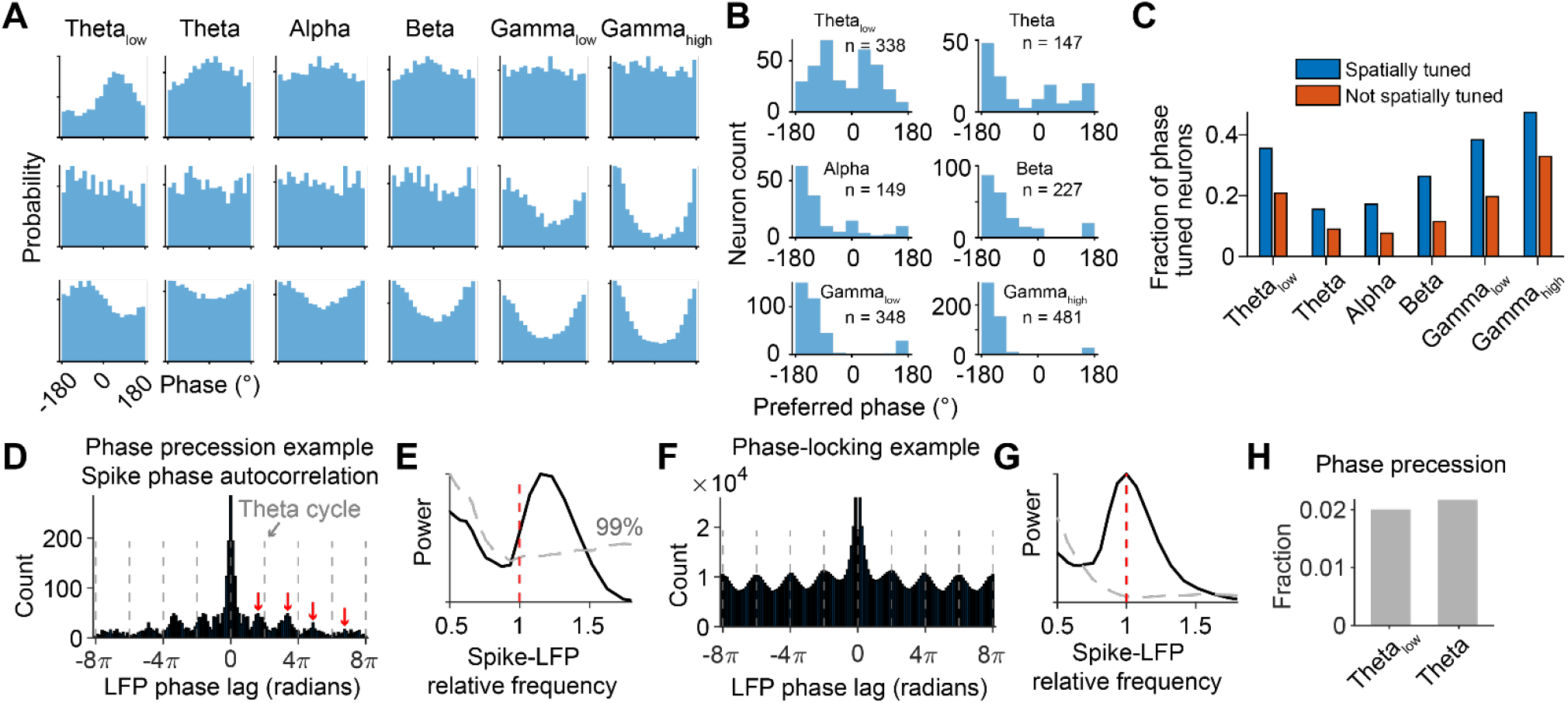
Spike-LFP phase coding. (A) Spike-LFP phase distribution for 6 frequency bands for 3 example neurons. (B) Distribution of preferred LFP phase for all significantly phase-locked neurons. (C) Bar plot of the fraction of phase-locked neurons in each frequency band for spatially-tuned and other neurons. (D) Spike phase autocorrelation for an example neuron. Dashed lines indicate theta cycle. Red arrows indicate peak autocorrelation in consecutive theta cycles. Note gradual advancement of spike phase relative to LFP phase. (E) Power spectrum of the autocorrelation shown in (D). Gray dashed line shows the 99% percentile of the shuffled distribution. Spike-LFP relative frequency larger than 1 indicates spikes oscillate at a higher frequency than theta. (F) and (G) The same as (D) and (E) for an example phase-locking neuron without phase precession. (H) Fraction of neurons showing significant phase precession for low theta and theta.

In the rodent hippocampus, place cells fire at progressively earlier theta phase within the place fields. We searched for this property in freely-moving macaques. We calculated the autocorrelogram of spike-LFP phases and computed how fast it oscillates relative to LFP phase: If spike-LFP phases oscillate at a higher frequency than LFP phase, it indicates phase precession (Mizuseki et al., 2009). We found that some neurons fired at gradually earlier theta phases in consecutive theta cycles (Fig. 6D), resulting in a faster spike-phase frequency relative to the corresponding LFP theta frequency (Fig. 6E) (Methods). In contrast, for phase-locking neurons, spike-phase frequency was the same as LFP frequency (Fig. 6F,G). Theta phase precession was only present in a handful of cells (12/599 neurons for low theta, 13/599 neurons for theta) (Fig. 6H), which were all tuned to various spatial variables.

These results suggest that spike-LFP phase coding is similar, but theta phase precession less prevalent in macaques compared to rodents.

### Facing location tuning reflects heading- but not viewing-related properties

Our FL variable is similar to ‘spatial view’ (SV) – where the monkey looks in the environment – described in head-fixed macaques (Georges-François et al., 1999; Rolls et al., 1997), but referenced to head rather than gaze. Combining eye-in-head movement and 3D head orientation allowed us to disambiguate between SV and FL (Fig. 7A). Only about 11% (28/244) of HF cells were significantly tuned to SV, as compared to 24% (58/244) for FL, when either variable was included in the model. Critically, including both SV and FL in the model revealed that neural activity was predominantly driven by FL rather than SV (46 vs. 8 tuned neurons; Fig. 7B). We performed a similar comparison for HD vs. gaze direction (GD) and found that tuning again reflected mostly head rather than gaze (15 vs. 5 tuned neurons; Fig. 7B).

**Fig. 7.**
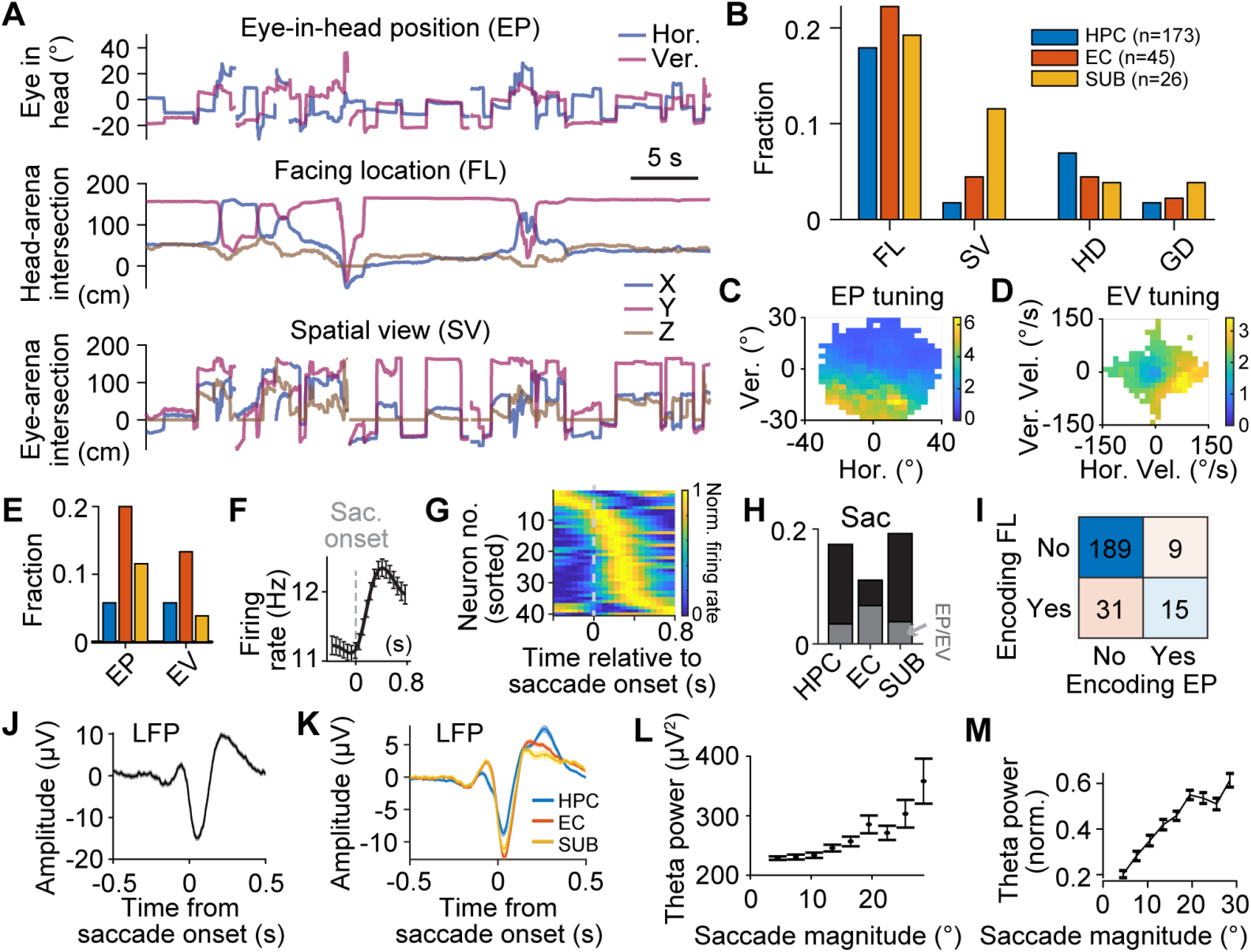
Allocentric facing location and head direction tuning predominantly reflects head but not gaze properties. (A) Example raw traces showing eye-in-head position (EP), facing location (FL), and spatial view (SV). (B) Fraction of neurons in each region encoding FL, SV, azimuth head direction (HD) and gaze direction (GD) when fitting all head- and gaze-related variables simultaneously. (C) Colormap representation of the firing as a function EP for an example tuned neuron. (D) Colormap representation of the firing as a function of eye-in-head velocity (EV) for an example tuned neuron. (E) Fraction of neurons in each region (color-coded bars, same as (B)) encoding EP and EV. (F) Mean firing rate from saccade onset for an example tuned neuron. Error bars, SEM over saccade events. (G) Colormap of the normalized firing rate for all saccade-tuned neurons, sorted by the peak activity relative to saccade onset. (H) Fraction of neurons in each region (x axis) encoding saccade event. Overlapping neurons encoding EP or EV are shown as stacked bars. (I) Confusion matrix showing the number of neurons encoding (or not) FL and EP. (J) Average saccade onset-triggered LFP for an example session. Shaded area, SEM over saccade events. (K) Average saccade onset-triggered LFP for all regions. Shaded area, SEM over sessions in individual regions. (L) Theta band power vs. saccade magnitude for an example session. Error bars, SEM over saccade events. (M) Average, normalized theta band power vs. saccade magnitude for all sessions. Error bars, SEM over sessions. Data from monkey K. See also **Fig. S5**.

A small proportion of EC cells have been reported to show grid-like tuning when head-fixed macaques visually explored novel images on a screen (Killian et al., 2012). We performed a similar analysis searching for grid tuning in the SV variable. Based on the same criterion used by Killian et al. (2012), 9% of EC neurons (4/45) showed grid-like tuning for SV as revealed using traditional grid cell analysis (Fig. S5E). But this effect was weak: no neurons showed grid tuning when we increased the significance level from 95% to 99% (Fig. S5E). There were no significant grid neurons (at 99%) also in HPC and SUB. Similar results were obtained for FL tuning: 7% of EC cells showed grid tuning at 95% but zero at 99% significance level.

We also examined egocentric eye (eye-in-head) tuning. Consistent with studies in head-fixed monkeys (Meister and Buffalo, 2018) and a recent study in mice (Mallory et al., 2021), we found neurons tuned to eye position (EP) and/or eye velocity (EV) (Fig. 7C,D), with the EC showing the strongest tuning (22%; Fig. 7E). HF neurons (16%) were also modulated by saccadic eye movements, forming saccade-locked sequential activity (Fig. 7F,G). Unlike EP/EV tuning, saccade tuning was stronger in the HPC/SUB than EC (Fig. 7H). We also examined mixed selectivity between the dominant spatial variable (FL) and EP: 15 neurons encoded both FL and EP, but coding of FL only was more common than EP only (Fig. 7I).

In primates, saccades synchronize network activity across regions (Sobotka and Ringo, 1997). We found that hippocampal LFP consistently showed a slight positive deflection, followed by a large negative deflection and another positive deflection that was linked to saccade events (Fig. 7J), resonating with a previous report (Jutras et al., 2013). The first positive and negative deflections were smaller, but the second positive deflection was larger in the HPC than the EC and SUB (all p < 0.001, Kruskal-Wallis test) (Fig. 7K). In addition, consistent with a previous study in head-fixed monkeys (Doucet et al., 2020), theta band power was positively correlated with saccade magnitude (Pearson’s r = 0.96, p < 0.001) (Fig. 7L,M). These results suggest that the relationship between theta activity and saccade in macaques may be analogous to that between theta and locomotion in rodents.

In summary, macaque hippocampal activity shows rich eye movement-related signals. Nevertheless, allocentric FL and HD tuning predominantly reflects head, but not gaze, properties.

## Discussion

These results provide a comprehensive picture of novel spatial coding schemes in the macaque HF, for the first-time during free exploration with accurate tracking of head motion and eye movement. At the network level, intermittent theta band activity was locked to movement onset. At the single cell level, head facing location, egocentric boundary and 3D head direction dominated over allocentric place coding, contrary to what is generally accepted from rodent studies. We also found negligible grid tuning in freely-moving macaques, and this was true regardless of whether it was computed relative to allocentric space, facing location or spatial view. Macaque hippocampal activity was also strongly modulated by eye movements - reported previously only in head-fixed studies. Nevertheless, spatial tuning of orientation variables was dominantly anchored to head, more than gaze.

### Behavioral correlates of theta activity and theta phase coding

A salient finding is the intermittent, lower frequency (~4 Hz) theta activity, restricted to movement onset in macaques, as compared to a continuous rhythm in locomoting rodents (~6-10 Hz) (Buzsáki and Moser, 2013). Nevertheless, like rodents, this low theta activity showed speed-dependent frequency increases. Human studies have also consistently reported low frequency oscillations around 3-4 Hz (Bush et al., 2017; Ekstrom et al., 2005; Watrous et al., 2011), but other studies have reported higher theta frequencies, particularly during real world navigation (Aghajan et al., 2017; Bohbot et al., 2017; Goyal et al., 2020). The fact that theta LFP has a different function between primates and rodents has also been reported by a recent macaque study (Talakoub et al., 2019), who showed that theta oscillations (i) were of low frequency (~4 Hz) and not sustained during walking behavior, and (ii) predicted sleep instead of active states.

Interestingly, despite low power theta oscillations in macaques, theta phase coding is nonetheless conserved: A significant fraction of neurons were phase-locked to low theta (56%) and theta (25%). Despite the prevalence of phase-locking, theta phase precession existed only in a very small number of HF neurons of freely-behaving macaques. This low incident of theta phase precession in the macaque HF contrasts with its abundance among rodent place cells (Mizuseki et al., 2009); notably, this property is also less prevalent for bat place cells (23%) (Eliav et al., 2018). It remains to be determined whether theta phase precession becomes more prominent when macaques engage in memory/decision tasks, as well as whether and how distinct theta bands and theta phase precession are tied to specific aspects of the task.

### Head facing and 3D head direction tuning

The finding that facing location and head tilt represent the dominant tuning variables stresses the importance of accurately tracking 3D head orientation during free behavior. Whereas facing location has been similarly described previously in macaques in the form of ‘spatial view’ cells (Rolls et al., 1997), the strong head tilt correlates across HF regions may appear surprising. Nevertheless, tilt tuning has been shown to be prevalent in the bat presubiculum (Finkelstein et al., 2015), the monkey anterior thalamus (Laurens et al., 2016) and the mouse anterior thalamus and retrosplenial cortex (Angelaki et al., 2020). Thus, orientation relative to gravity may be an important component in spatial coding and behavior across species.

It is currently unknown whether rodent HF cells encode facing location. The strong prevalence of facing location/spatial view tuning in monkeys, along with the absence of rodent-like robust place fields, represents the most notable difference in HF coding between the two species. Note that we could not distinguish whether the dominant variable encoded in the macaque HF truly represents facing location on the walls and ceilings of the open arena or, alternatively *egocentric head direction* relative to environmental landmarks (i.e., any salient cue in the environment, be it objects, boundaries, goals, etc.). We defined FL in a way analogous to ‘spatial view’ (Rolls et al., 1997), although we note that what this variable represents exactly must be further explored in future studies.

### Eye movement encoding

Previous studies in which the animal’s head was fixed in a cart failed to disambiguate the influences of head and eye movement on hippocampal responses (Georges-François et al., 1999; Rolls et al., 1997). When FL was substituted by SV, we found that some cells appear to encode where the eye looks. However, considering both variables together solved this ambiguity – FL of the head outperformed SV of gaze. Similarly, the firing of most direction-tuned neurons was better accounted for by head, rather than gaze. We propose that, because primates saccade continuously to explore their environment, head-anchoring may provide a more stable representation.

More important than the debate between FL vs. SV is the presence of eye movement signals in the macaque HF. Many neurons, particularly in the EC (22%), were tuned to egocentric eye-in-head position and/or velocity. This number is similar to results obtained in head-fixed monkeys viewing images (11.9% Grid + 9.3% Border in Killian et al., 2012; 20.3% in Meister and Buffalo, 2018; 22% in the HPC in Nowicka and Ringo, 2000). In addition, many cells, particularly in the HPC and SUB, were modulated by saccadic eye movements. Furthermore, head-fixed studies have found hippocampal LFP synchronization and amplitude modulation by saccades (Doucet et al., 2020; Jutras et al., 2013). We extended these findings to freely-behaving monkeys and further showed differential effects across HF regions. These results directly link eye movements with the navigation/memory circuit during natural behavior.

Despite strong eye movement-related responses in the HF, we found that the macaque EC showed little grid tuning both for allocentric location and for where the animal is looking in the environment. Previously, a small percentage (11.9%) of EC cells were reported to show grid tuning based on eye movements in head-fixed macaques (Killian et al., 2012). When we used a 95% criterion, as the authors did, we found 9% of EC neurons that passed the criterion for grid tuning. However, the tuning was so weak that it failed to reach significance at the 99% level for all neurons. Thus, visual grid tuning, even if present in a small number of EC cells, is far less robust, and possibly insignificant, compared to rodent grid cells. Alternatively, visual grid tuning may only be seen in head-fixed animals exploring a visual image and not in freely foraging primates. Future studies should directly compare visual space grid tuning between head-fixed virtual navigation (or viewing a steady image) and free navigation in open environments on a cell-by-cell basis.

Combining present and previous findings, head facing may act as an anchor for the firing fields in the HF, whereas eye movements may be used to establish a more fine-grained map for things seen centered around that facing location. The eyes actively interrogate and efficiently gather information by swiftly parsing complex scenes. Subtle eye movements may reflect internal deliberation and the prioritization of goals in real-time (Gottlieb and Oudeyer, 2018; Yang et al., 2016). The reported eye movement encoding in spatial navigation circuits may also reflect task-relevant variables that covary with eye movements. It is well known, for example, that eye movements both guide and offer a window into cognition and memory, which could be mediated by the HF (Hayhoe and Ballard, 2005; Nau et al., 2018; Ryan and Shen, 2020). Eye movements have also been linked to infer subjects’ beliefs by continuously tracking an invisible target (Lakshminarasimhan et al., 2020). Accurate goal-tracking by gaze was associated with improved task performance, and inhibiting eye movements impaired navigation precision. Thus, it is important that future experiments explore the precise advantage that active visual sampling confers upon navigational planning and the extent to which humans and monkeys exploit it.

### Position (‘place’) coding in freely-moving monkeys

Since the discovery of place cells in rodents, it has been a long-standing question whether the primate HF represents self-location in a similar way during free behavior. It is important to note that, using traditional tuning curve analysis, the proportion of place coding cells in the HPC is not different from previous monkey studies (present study: 26% in HPC, as compared to 30.6% in Matsumura et al., 1999; 25.4% in Hazama and Tamura, 2019; 32.1% in Ludvig et al., 2004 - squirrel monkeys; 14.1% in Courellis et al., 2019 - marmosets). Further, in addition to their low occurrence compared to rodents, ‘place fields’ have been reported to be more dispersed in moving marmosets than rodents (Courellis et al., 2019), a finding that is consistent with the present results. Thus, the difference in conclusions is not data-driven, but rather analysis-driven. Nevertheless, we note that we cannot exclude the possibility that the lower fraction of place and grid coding is due to the restricted size of the arena or recording location; strong place and grid coding may be found in locations/cortical layers not sufficiently sampled in the present recordings. In the wild, the natural habitat of rhesus macaques is much larger than the arena used in the present study, but our monkeys have always lived in small environments.

Importantly, unlike previous studies, when accounting for other variables quantitatively using a multi-modal model, the percentage of true place cells dropped from 26% to 7% in the HPC, a percentage that is lower but still significant. This difference emphasizes the need to avoid the use of traditional tuning curve analysis in neurons with mixed selectivity. Although the vertical coverage is limited in the present study, a sizable proportion (13%/19%) of HPC/EC cells are tuned to either horizontal or vertical head position. Thus, like head direction, place coding in freely-moving macaques is three-dimensional, as previously shown for both rats and bats (Grieves et al., 2020; Yartsev and Ulanovsky, 2013). The present study found a negligible proportion (2%) of grid coding. We note that, allocentric border cells would also show as position-coding; thus such cells are also infrequently encountered in the macaque HF.

Putative place cells have been reported in human patients during virtual navigation (26% in Ekstrom et al., 2003; 25.6% in Miller et al., 2013). Single cell recordings have also confirmed grid-like tuning in the human EC in a virtual navigation task (14% in Jacobs et al., 2013; 18.4% in Nadasdy et al., 2017). Indirect measurements in fMRI studies have further corroborated grid-like activity in the human EC, that is also modulated by speed (Doeller et al., 2010). In a recent human study during a spatial memory task, hippocampal neurons showed selectivity to remote goal location, more so than self-location, with some cells also encoding heading direction and trial progression (Tsitsiklis et al., 2020). Future experiments should compare and contrast HPC/EC spatial representations during goal-oriented and free foraging.

### Topography of spatial coding in hippocampal-neocortical networks

In primates, the posterior hippocampus is analogous to the dorsal hippocampus in rodents (Strange et al., 2014). The posterior hippocampus showed stronger spatial tuning to facing location, head tilt, and egocentric boundary than the anterior hippocampus. The posterior hippocampus is also highly connected with the posterior neocortex, including retrosplenial and parietal cortices, which are strongly linked to spatial processing (Clark et al., 2018). Therefore, the topographical differentiation of spatial tuning may reflect one aspect of a general organization of the hippocampal-neocortical networks in different aspects of cognition, in that the posterior part is more involved in detailed spatial representation and the anterior part is more involved in emotion-related processing (although this still remains controversial) (Strange et al., 2014). The technical innovations will now allow us to interrogate how different cognitive aspects segregate and interact across extended hippocampal-neocortical networks using large-scale wireless recordings in freely-moving primates.

## Acknowledgments

We thank J. Lin and B. Kim for help with setting up the arena; G. DeAngelis for help with surgeries; M. Schartner for help with the video-based tracking system; N. Tataryn for veterinary care; L. Lu and K. Bohne for advice on the tetrode technique; B. Goodell and C. Gray for assistance with microdrive design; K. Lakshminarasimhan for help with the model fitting.

## Author contributions

D.M. and D.E.A. designed the study. D.M., E.A., and B.C. set up the arena. D.M., E.A., and J.D.D. performed the surgeries. D.M. performed the experiments and analyzed the data. J.L. contributed to the analysis. D.M. and D.E.A. wrote the manuscript with inputs from all other authors.

## Data and materials availability

All data and code are available from the corresponding author upon request.

## Supplemental information

### Materials and Methods

#### Animals

Three male rhesus macaques (*Macaca mulatta*, 8-10 years old) weighing 9-14 kg were used in this study. Animals were chronically implanted with a lightweight polyacetal head ring for head restraint (Meng et al., 2005). A head cap printed with carbon-fiber reinforced nylon (Utah Trikes, USA) was attached on the head ring to accommodate and protect the microdrives, recording devices, transmitters, and batteries. We used the standard pole-and-collar method to train the monkeys to move from their home cage to the primate chair using positive reinforcement. All animal experimental procedures and surgeries were approved by the Institutional Animal Care and Use Committee at Baylor College of Medicine and were in accordance with the National Institutes of Health guidelines.

Pre-op computed tomography (CT) and magnetic resonance imaging (MRI; 3T, Siemens) images were collected to segment the regions of interest, and for accurate design of the form-fitting (to the skull) chambers (titanium) and the microdrives (form fitting to the brain surface; Visijet-Clear or Ultem). CT and MRI images were registered in 3D slicer software (Fedorov et al., 2012) (https://www.slicer.org). The hippocampus, entorhinal cortex, subicular complex, and the entire brain were segmented semi-manually from the T1-weighted MRI images in ITK-SNAP software (Yushkevich et al., 2006) (www.itksnap.org), with reference to standard atlas (Paxinos et al., 2008; Saleem and Logothetis, 2006). Major blood vessels were segmented from the T2-weighted MRI images. The skull model was segmented from the CT images. All rendered 3D models were exported from ITK-SNAP and then imported into 3D slicer, where MRI and CT volumes were aligned with stereotaxic coordinates. Models of the chambers and microdrives were imported in 3D slicer and the best positioning was chosen such that the electrodes covered the largest extent of the hippocampus and the posterior entorhinal cortex (homolog to the medial entorhinal cortex in rodents). Major blood vessels were avoided.

#### Freely moving monkey (FMM) arena

The FMM arena consisted of an open circular enclosure with a 3.30 m diameter and a 2.12 m height, with a single entrance/exit door and a drain on the floor (Figure S2). The enclosure was made of white composite material. Three feed/touch-screen boxes were evenly located at the perimeter of the arena. A motion capture system (Vicon Ltd, UK) consisting of 9 infrared cameras (Bonita camera) was mounted in the ceiling and the wall (Figure S2), to capture the 3D position of the 4 reflective markers placed on the monkey’s head cap at a rate of ~1 kHz. Nine video cameras (CM3-U3-13Y3M 1/2” Chameleon3 Monochrome Camera, FLIR Systems, Inc., USA) were mounted on the wall to capture videos at ~30 Hz from different angles through a transparent plastic window. A wide-angle camera was mounted in the center of the ceiling for surveillance. The arena was lit by an LED lamp mounted at the ceiling.

#### Implantation of chronic microdrives

We used two different technologies for recordings.

First, Monkeys B and L were implanted with the NLX-18 and NLX-9 tetrode drives (Neuralynx, Inc., USA) using a tetrode-in-guide tube technique. First, guide tubes (2” long 27G needles) were loaded and fixed inside a grid, which rested inside a chamber. The depths of the guide tubes were planned such that they were positioned ~3-5 mm above the target regions. Second, a polymicro capillary tubing (100 μm ID, 170 μm OD; Molex, LLC, USA) was loaded into the guide tubes and glued to the drive shuttles. Third, each tetrode (Platinum 10% Iridium, 0.0007”; California Fine Wire Co., USA) was loaded into the polymicro capillary tubing. The tetrode was glued to the capillary tubing and its tip extended outside from the bottom of the tubing by ~2-3 mm. The NLX-18 drive was loaded with 16 tetrodes and the NLX-9 with 8 tetrodes. The tetrodes were plated with platinum to an impedance of 100-200 kΩ. The entire drives were gas-sterilized (EtO) before surgery. Monkey B received an NLX-18 drive in the right hemisphere. Monkey L received an NLX-18 drive in the right hemisphere and an NLX-9 drive in the left hemisphere. The assembled drive was implanted as a single unit and only a single surgery was required.

Second, Monkey K was implanted with a 32-channel microdrive (SC32) in the left hemisphere and a 124-channel microdrive (LS124; Gray Matter Research, LLC, USA) in the right hemisphere (Dotson et al., 2017). Each channel was loaded with a glass-coated tungsten electrode (250 μm total diameter, 60 ° taper angle, ~1 MΩ impedance; Alpha Omega Co., USA). Each electrode had a travel distance of 42 mm from the brain surface. The implantation procedure breaks into 3 separate surgeries (Gray Matter Research) (https://www.graymatter-research.com/documentation-manuals/): Chamber implantation, craniotomy, and microdrive implantation. In stage 1, the chamber was implanted at the desired location using C&B Metabond (Parkell, Inc., USA) and dental cement; bone anchor screws were not required since everything was protected within the head cap. The chamber was hermetically sealed with a plug and O-rings. At 10 days after stage 1, we collected fluid sample from the inside of the chamber and cultured the sample. After no signs of infection and a negative culture, we moved on to stage 2. In stage 2, a craniotomy was made inside the chamber and the edge of the craniotomy was polished with 90 ° Kerrison rongeurs. A form-fitting plug (with the same shape as the microdrive) was installed and the plug & chamber unit were hermetically sealed. At 10 days after stage 2, we controlled for infection again by collecting and culturing a fluid sample from the inside of the chamber. After no signs of infection and a negative culture, we moved on to the final stage. In stage 3, the microdrive, loaded with electrodes, was inserted into the chamber and secured to the chamber walls using multiple screws. The entire assembly was then hermetically sealed with O-rings. All components were autoclaved or gas-sterilized prior to procedures.

For Monkeys B and L, all tetrodes were advanced to 1 mm out of the guide tubes on the day of implantation. For Monkey K, all electrodes targeting deep structures were moved to 8 mm below brain surface on the day of microdrive implantation and advanced to a position of 2-3 mm above target areas within 5 days following recovery from surgery. Electrodes were advanced by turning fine threaded screws, at a resolution of 125 μm per revolution for single electrodes and 250 μm per revolution for tetrodes. Electrodes were advanced by 50-250 μm per day and recording was performed the next day. A CT scan was performed every 2-3 months to reconstruct current electrode locations. Electrodes were visible as white tracks in the CT images (Figures 1 and S1). CT volumes were registered with pre-op MRI volumes to visualize electrode locations in the brain (Premereur et al., 2020). When combined with electrode advance history, electrode locations were reconstructed with good precision. Anatomical landmarks such as ventricles, white matter and electrophysiological signatures were used to further verify electrode locations.

We additionally verified electrode locations with histology in Monkey B. The monkey was perfused with electrodes in position. After post-perfusion fixation, the brain was removed and sunk in fixative with 30% sucrose for several days. The brain was sectioned at 60 μm and stained with cresyl violet.

#### Behavioral training and tracking

Monkeys were placed on a food delayed schedule. They were fed fewer than normal biscuits (~15%) in the morning so they were encouraged to forage for treats in the arena. The remaining daily allotment of food less the amount received in the arena was fed after the monkeys were returned to the home cage after training, and supplemented with vegetables and fruits. The monkeys were habituated to the FMM arena in a gradual manner. They were brought from their home cage to the arena in a transfer cage. They entered/exited the arena through a single door (same one each time). They were trained to freely forage for randomly scattered food pellets or fruits on the floor throughout the session. Monkeys B and L also received rewards from the reward boxes with equal probability. Monkey K did not forage at the reward boxes. At the end of each session, the monkeys were allowed to return to the transfer cage and then back to the home cage, where they received food and additional treats. Monkey K was habituated to wear a wireless eye tracking device (ISCAN, Inc., USA). The monkey was initially trained to wear a dummy eye tracking device, which could be replaced inexpensively when destroyed, in the home cage. Once the monkey stopped interacting with it, they were trained to wear the eye tracking device in the FMM arena. The eye tracking device was housed in a rigid, 3D-printed case which was fixed to the head ring implant. The device consists of a miniature infrared camera, an infrared emitter, a hot mirror, and a transmitter. The hot mirror was held rigidly in front of the left eye and the camera faced downward at about 45 ° relative to the mirror.

To accurately track monkey’s head motion in 3D, we used a marker-based motion tracking system. Four markers were placed within a single plane on the head cap (Figure 1H). One marker was placed in the front and one at the back. Other two markers were symmetrically placed on the left and right (closer to the back marker). The 3D position (x,y,z) of each marker was recorded (~1 kHz) in Spike2 with a data acquisition system (Power1401, CED Ltd., UK). The eye tracking device was powered by a lightweight 3.7 V Li-Po battery. Eye images were wirelessly transmitted to a receiver outside the arena. Horizontal/vertical eye position was monitored (~30 Hz) in Spike2.

#### Electrophysiological recordings

We used a 64-channel neural logger to record broadband (0.1-7000 Hz) electrophysiological signals at 32 kHz (Deuteron Technologies Ltd., Israel). The neural logger was powered by a lightweight 3.7 V Li-Po battery. The head-stage was connected to the drive connectors directly or via a 5 cm jumper cable. The head-stage utilized a preamplifier and a 16 bit analog-to-digital converter from Intan Technologies. The signal precision was 0.2 μV. Raw signals were digitized on board and stored on a 64 GB micro-SD card (SanDisk) plugged into the logger. To avoid clock drift over time, the logger was wirelessly synchronized with the computer clock every 5-10 minutes via a USB transceiver placed outside the arena; the transceiver was connected to the Spike2 system for synchronization between behavioral data and electrophysiological recordings. Each recording session usually lasted ~20-60 minutes. For Monkey K, the chamber was used as the ground. For Monkeys B and L, one guide tube or a separate screw held in the skull was used as the ground.

### Data analysis

All data analysis was performed in MATLAB 2019b (The MathWorks, Inc., USA).

#### Extraction of behavioral variables

Three-dimensional behavioral variables were extracted from the raw marker position data that were pre-processed to fill small gaps (< 1 s) which could happen occasionally. All raw data were re-sampled at 50 Hz.

Head-in-world *<u>position</u>* in the horizontal plane was defined as the *x* and *y* components of the average across all 4 markers. The amplitude of its time derivate was translational *<u>speed</u>. <u>Head height</u>* was defined as the *z* component of 3D position (arena floor at z = 0). *<u>Head tilt</u>* was defined as the projection of the earth gravity vector onto the head coordinate system; only pitch and roll angles (as a 2D variable) were considered. *<u>Azimuth head direction</u>* was calculated as in (Angelaki et al., 2020). Heat tilt and azimuth head direction, together, define 3D orientation. We computed *<u>angular velocity</u>* about the head’s yaw, pitch and roll axes. We focused on the yaw and pitch angular velocities (as a 2D variable) since the head rotated mostly along these 2 dimensions. *<u>Egocentric boundary</u>* was defined in a polar coordinate system, with the radial component calculated as the distance from the head center to the nearest arena boundary and polar angle defined between the arena center-to-head center vector and the azimuth head direction. To calculate where the monkey looked, we obtained 3D gaze direction by combining eye-in-head position and 3D head direction. The inner surface of the arena was treated as an enclosed cylinder, composed of three surfaces: the ceiling, the floor, and the cylinder wall. *<u>Spatial view</u>* was then computed as the intersection between the 3D gaze direction and these surfaces. *<u>Facing location</u>* was computed similarly, i.e., as the intersection between the 3D head direction and the three surfaces.

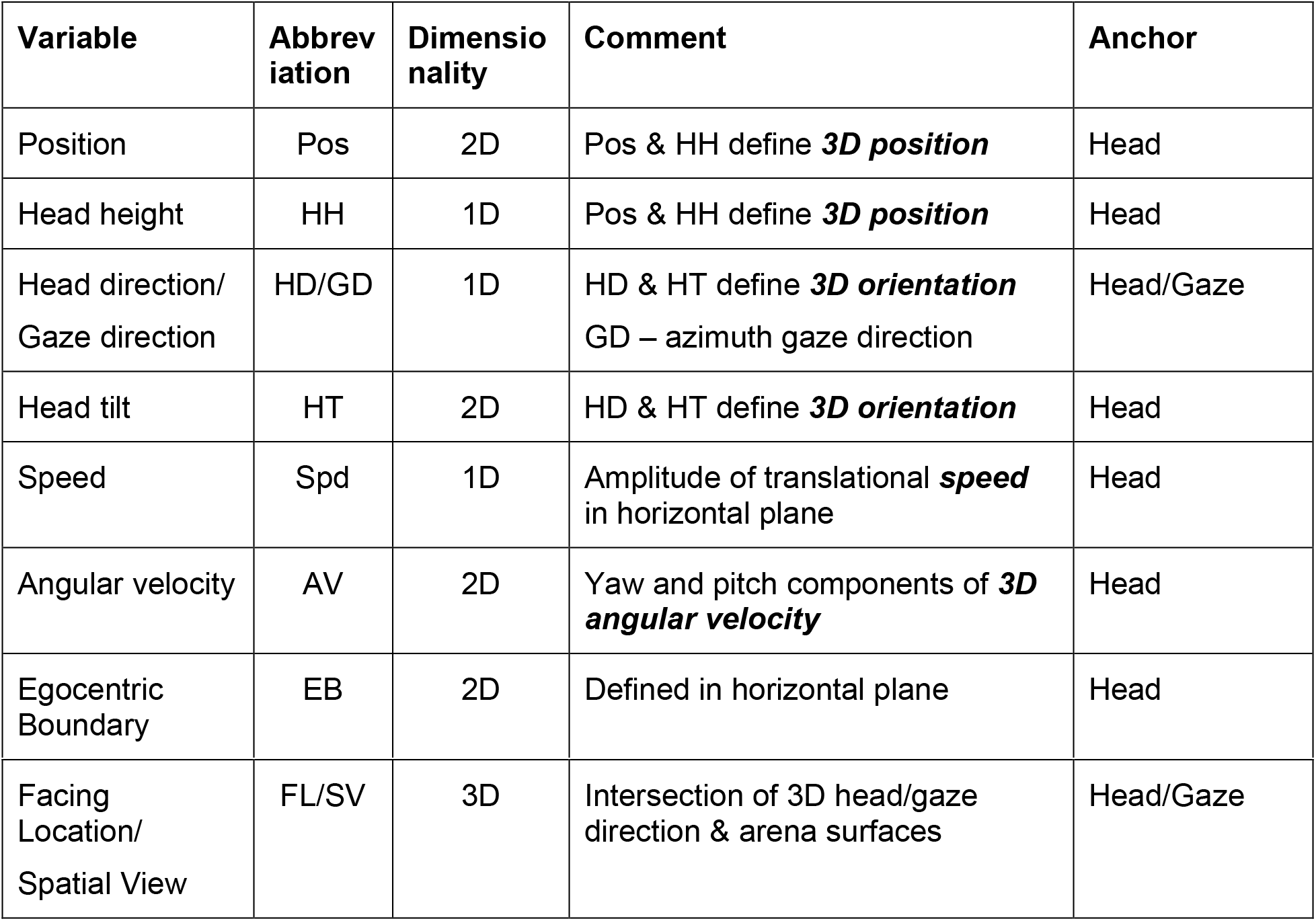

| Variable | Abbreviation | Dimensionality | Comment | Anchor |
| --- | --- | --- | --- | --- |
| Position | Pos | 2D | Pos & HH define **3D position** | Head |
| Head height | HH | 1D | Pos & HH define **3D position** | Head |
| Head direction/<br>Gaze direction | HD/GD | 1D | HD & HT define **3D orientation**<br>GD – azimuth gaze direction | Head/Gaze |
| Head tilt | HT | 2D | HD & HT define **3D orientation** | Head |
| Speed | Spd | 1D | Amplitude of translational **speed**<br>in horizontal plane | Head |
| Angular velocity | AV | 2D | Yaw and pitch components of **3D angular velocity** | Head |
| Egocentric Boundary | EB | 2D | Defined in horizontal plane | Head |

| Facing Location/<br>Spatial View | FL/SV | 3D | Intersection of 3D head/gaze direction & arena surfaces | Head/Gaze |
| --- | --- | --- | --- | --- |

For the model fitting, these behavioral variables were binned as follows: position (2D variable, −165-165 cm, 20 x 20 bins); facing location and spatial view (3D variable, x & y: −165-165 cm, z: 0-212 cm, 17 x 17 bins for the ceiling and floor, 53 x 11 bins for the wall); egocentric boundary (2D variable, x & y: −165-165 cm, 20 x 20 bins); head tilt (2D variable, pitch: −55-90 ° (up-down), roll: −18-18 ° (left-right), 18 x 6 bins); angular velocity (2D variable, −120-120 °/s, 16 x 16 bins); head height (1D variable, 10-70 cm, 12 bins); translational speed (1D variable, 0-120 cm/s, 15 bins); azimuth head direction (1D circular variable, −180-180 °, 18 bins). Sessions in which the monkeys occupied fewer than 80% of the valid position bins were excluded from further analysis.

Note that in this analysis, allocentric position-dependent spatial variables like border and grid tuning, will not be distinguished from Pos tuning. Thus, Pos tuning encompasses not only place, but also grid and border fields.

#### Preprocessing of electrophysiological data

We used a semi-automatic procedure for the preprocessing of the electrophysiological data. To obtain the LFP, the raw data were low-pass filtered at 300 Hz and down-sampled to 1 kHz. We used established pipelines, using the software Klusta for spike detection and automatic spike sorting and phy Kwik-GUI for manual curation (Rossant et al., 2016). Manual corrections were based on spike waveforms, waveform features, auto-correlograms, and cross-correlograms. Only well-isolated units were kept for further analysis. We used two independent measures to quantify spike sorting quality. In the first measure, we calculated the percent of spikes that fall within the refractory period (3 ms), i.e. refractory period violation. In the second measure, we calculated the isolation distance as the n^th^ smallest Mahalanobis distance of the spikes in other clusters from the current cluster in the feature space (n is the number of spikes in the current cluster) (Schmitzer-Torbert et al., 2005). For this measure to work, it must contain other clusters that have larger than n total spikes.

#### LNP model fitting

Several recent studies have emphasized the importance of using multimodal models to characterize mixed selectivity, which are agnostic to tuning curve shape and robust to the interdependence of encoded variables (Hardcastle et al., 2017; Laurens et al., 2019). Specifically, by comparing a multivariate linear-nonlinear Poisson (LNP) model framework with traditional classification methods based on tuning curves, Laurens et al. (2019) found that traditional analysis approaches often produce biased results. A fundamental advantage of the multivariate LNP model framework is that it can, by design, correctly identify multimodal responses even when various behavioral variables are correlated or interdependent. For example, position and egocentric boundary (EB) variables are typically strongly correlated, such that an EB-tuned cell may appear to be tuned as a place cell (Laurens et al., 2019).

Thus, to quantify the encoding of the behavioral variables by neuronal activity, we adapted this LNP framework, which has been successfully used in investigating the neuronal encoding of navigational variables in the rodent hippocampal formation (Hardcastle et al., 2017; Laurens et al., 2019; Ledergerber et al., 2020). The MATLAB code is available on GitHub (https://github.com/kaushik-l/neuroGAM). Spike trains were binned into 0.02 s time bins, the same as the behavioral variables. To test for significance of the model fitting and to control for overfitting, we used a 5-fold cross-validation approach. To obtain the training and test sets with minimal bias, the data were divided into 3 chunks; in each chunk, the data were further divided into 5 sub-chunks; for each fold during the cross-validation process, the *I^th^* (*i* ∈ (*1,2,3,4,5*), 5 folds) sub-chunk in each chunk was concatenated and used as the test set (20% of data) and the remaining used as the training set (80% of data). The measured spike train *r* was smoothed with a Gaussian kernel with 0.06 s (3 time bins) standard deviation. The best-fit parameter was solved using the MATLAB fminunc function (using the ‘trust-region’ algorithm). We used log-likelihood as the quantification for model performance (i.e. goodness of fit). The model performance was compared to a null model (mean firing rate model) to test significance (one-sided Wilcoxon signed rank test, alpha = 0.05).

We used an optimized forward search approach to select the best, simplest model. First, we fitted the 1^st^ order models that contained only one variable. Second, if any of the 1^st^ order models performed better than the null model, we fitted the 2^nd^ order models (2 variables) which included the best 1^st^ model variable (with the highest, significant log-likelihood value). Higher order models containing the best, significant lower model variables were fitted further until the model performance did not improve. The best simplest model was selected as the final best model.

The neuron categorization (Figure 4D) was based on the best 1^st^ order model; for example, if the 1^st^ order model containing the position variable had the best significant performance, this neuron would be categorized as a ‘position’ cell. The fraction of neurons encoding each variable (Figure 4E) was based on the final best model, in which a single neuron could encode more than 1 variable; for example, if the best model included egocentric boundary + translational speed + head direction, then this neuron is counted as encoding all these 3 variables. This approach avoided assigning variables that were significant in the 1^st^ order models but were not picked up by the final best model since the behavioral variables could be interdependent, which is usually not taken into consideration in traditional analyses.

To distinguish between the tuning to head and eye-related properties, in Fig. 7, we repeated the LNP model fitting by adding 2 eye-related variables: SV and GD, to the above model. Therefore, the final model included 10 variables (see the table above).

#### LFP analysis

We used the continuous wavelet transform (CWT) to compute the scalogram (Torrence and Compo, 1998). We used a model-based approach to compute the aperiodic as well as the oscillatory components in the power spectrum (https://github.com/fooof-tools/fooof). We extracted the average power in 6 frequency bands from the scalogram: low theta 1-4 Hz, theta 4-8 Hz, alpha 8-12 Hz, beta 12-30 Hz, low gamma 30-60 Hz, and high gamma 60-120 Hz. To extract LFP phase, we first band-pass filtered the raw LFP to individual frequency bands using a 4^th^-order Chebyshev Type II filter. We then used the Hilbert transform to extract instantaneous LFP phase. We used nearest neighbor interpolation to identify the corresponding LFP phase for each spike. We used Rayleigh test to identify significant spike-LFP phase tuning with alpha = 0.01.

To analyze LFP phase precession, LFP phase in the theta bands was unwrapped such that the resulting phase is monotonic instead of cyclic (Mizuseki et al., 2009; Qasim et al., 2020). For each spike, we assigned the corresponding unwrapped LFP phase via nearest neighbor interpolation. We excluded spikes where the filtered LFP amplitude fell in the lowest 10 percentile. We calculated the autocorrelogram (22.5° bin) of the spike-LFP phase trains. We then calculated the power spectrum of the autocorrelogram. If a neuron is phase-locked to LFP phase, the power spectrum peaks at the same frequency of the autocorrelogram relative to the LFP phase (relative frequency = 1). If the power spectrum peaks at a faster frequency of the autocorrelogram relative to the LFP phase (relative frequency > 1), it indicates that spikes move to earlier LFP phases (i.e. phase precession). To identify significant phase precession, we assigned each spike a random, unwrapped LFP phase within the same theta cycle, and repeated the analysis for 500 times. We identified significant phase precession if the actual vs. shuffled power spectrum shows a significant peak in the range of relative frequency of 1 to 1.5 (alpha = 0.01/26 bins, corrected for multiple comparisons).

#### Circular graph for conjunctive coding

The degree of conjunctive coding was defined as the similarity of the encoding between variables, i.e. the size of the intersection divided by the size of the union – Jaccard index. The circular graph in Figure 4G included only variable pairs which had a Jaccard index above 0.1 (code at https://github.com/paul-kassebaum-mathworks/circularGraph).

#### Tuning curve clustering analysis

Tuning curve clustering analysis (Figures 5G and 5H right) was based on the model-based tuning curves. We first normalized the tuning curves so that the firing rate ranged from 0 to 1 then performed principal component analysis on the correlation matrix of all tuning curves. The first two principal components were plotted (example code at https://github.com/PeyracheLab/Class-Decoding). Uniformity test for data in Figure 5H used Rayleigh test in the CircStat toolbox (Berens, 2009) (http://bethgelab.org/software/circstat/).

#### Traditional tuning curve-based analysis

The model-based, unbiased approach employed in the present study is ideal for disambiguating the contributions of many variables, compared to more traditional analyses focusing on a single or limited number of variables, which can often over-represent the coding of variables considered. Nevertheless, to be able to compare with previous studies, we also performed a more traditional tuning analysis considering the most commonly used spatial variables.

Specifically, we used traditional methods to identify place tuning based on spatial information (*SI*) (Skaggs et al., 1993). We used grid score (*GS*) to quantify grid tuning (Langston et al., 2010). We set a speed threshold of 10 cm/s below which spikes were not considered. We first constructed the position activity map (firing rate as a function of position) using 6×6 cm binning (55×55 bins for x and y), which was then smoothed using a 2D Gaussian kernel with 9 cm standard deviation along each dimension. We used speed modulation depth to identify neurons tuned to speed. We first constructed speed tuning curves (firing rate as a function of speed, 30 speed bins, from 0-120 cm/s, 6 cm/s standard deviation Gaussian smoothing). We calculated speed modulation depth as the difference between the maximum and minimum speed tuning firing rate (Kropff et al., 2015). We used mean vector length to quantify azimuth head direction tuning. We first constructed circular head direction tuning curves (36 bins, 10°**/**bin, 15° standard deviation Gaussian smoothing). The mean vector length was calculated as the weighted sum of the tuning curve in each direction divided by the total firing rate across all directions using the CircStat toolbox (Berens, 2009). For grid analysis of the SV and FL variables, we computed the SV and FL activity map on the floor using 110×110 bins, smoothed with a 2D Gaussian kernel with standard deviation of 2×2 bins; and SV and FL activity map on the wall using 500×100 bins (perimeter x height), smoothed with a 2D Gaussian kernel with standard deviation of 5×1 bins along each dimension. We then computed *GS* to quantify SV and FL grid tuning either on the wall or on the floor.

To obtain the shuffled distributions, we circularly shifted the spike trains relative to the behavioral data for a random interval between 10 seconds and session duration minus 10 seconds, and re-calculated each measure mentioned above. We repeated this process 500 times. If the raw measure was larger than the 99^th^ percentile of the shuffled distribution, the neuron was considered tuned to the specific variables. We additionally set a *SI* threshold of 1 bits/spike to identify neuron tuned to position.

#### Eye movement analysis

We used a speed threshold of 150°/s to detect saccade events in either horizontal or vertical eye movements. We computed the slow phase eye velocity by taking the derivative of smoothed eye position (excluding saccade events). We fitted eye-in-head position (2D variable, horizontal: −40-40°, vertical: −30-30°, 20 x 20 bins) and velocity (2D variable, horizontal/vertical: −150-150°/s, 20 x 20 bins) with the LNP model (single variable model) to identify neurons tuned to either of these two variables. To identify neurons that were tuned to saccade events, we identified neurons that showed a significant difference between pre- and post-saccadic activity (0.4 s before and 0.4 s after saccade onset; paired *t*-test, alpha = 0.01).

All p values smaller than 0.001 were indicated as p < 0.001; otherwise exact p values were indicated.

---

**Figure S1.**
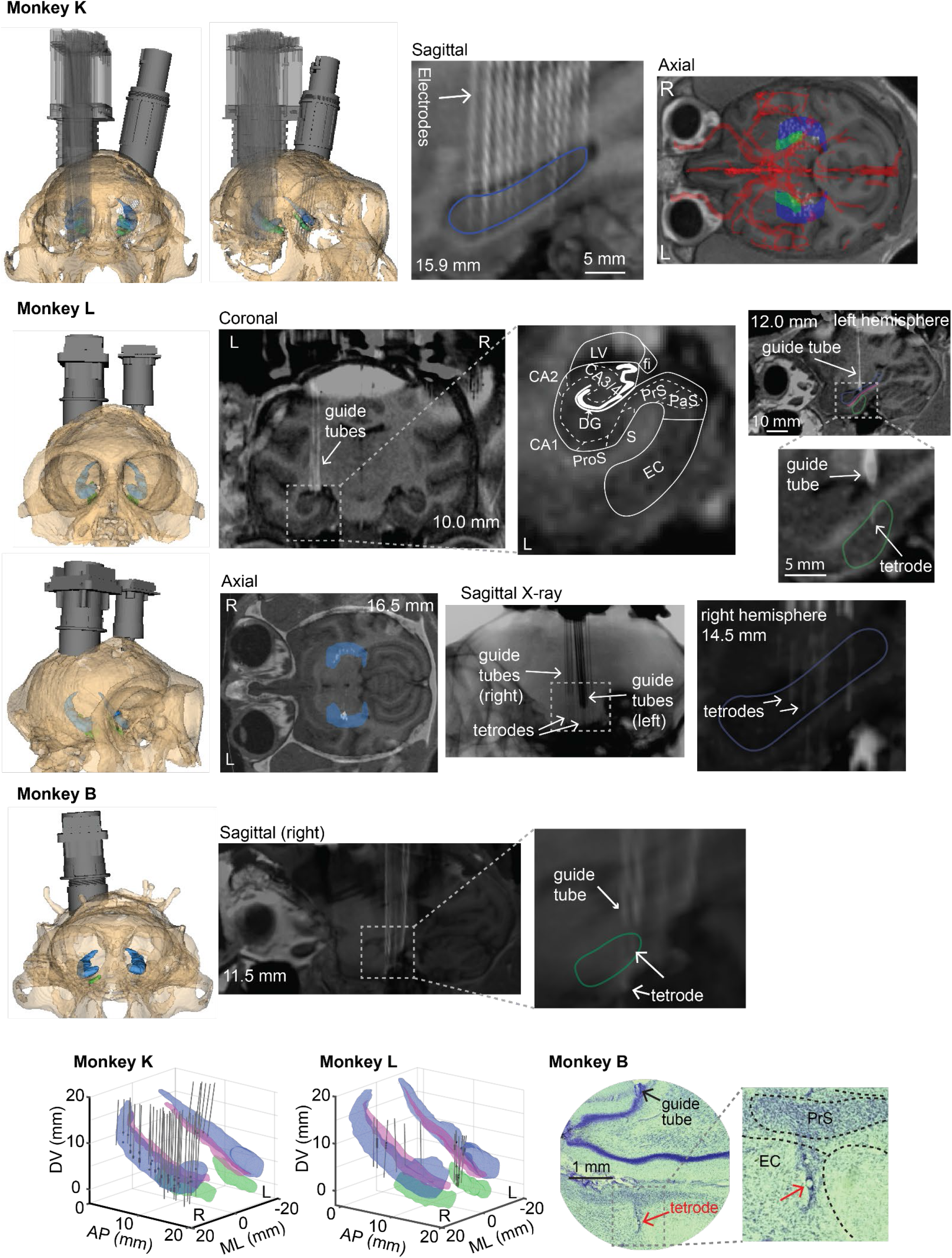
Microdrive implants and reconstruction of electrode locations, related to Figure 1. Microdrive (gray) and skull (bone white) models are shown for all three monkeys. Co-registered MRI and CT images show electrode locations as white tracks or dots. Hippocampus (HPC): blue, including CA1, CA2, CA3/4, and DG; Entorhinal Cortex (EC): green; Subicular complex (SUB): magenta, including subiculum (S), presubiculum (PrS), parasubiculum (PaS), and prosubiculum (ProS). Tetrodes were used for monkeys L and B; single electrodes were used for monkey K. Bottom, reconstructed models of HPC, EC, SUB and electrodes (gray lines). Locations where single units were recorded are indicated as black dots. An example histology image is shown for monkey B.

**Figure S2.**
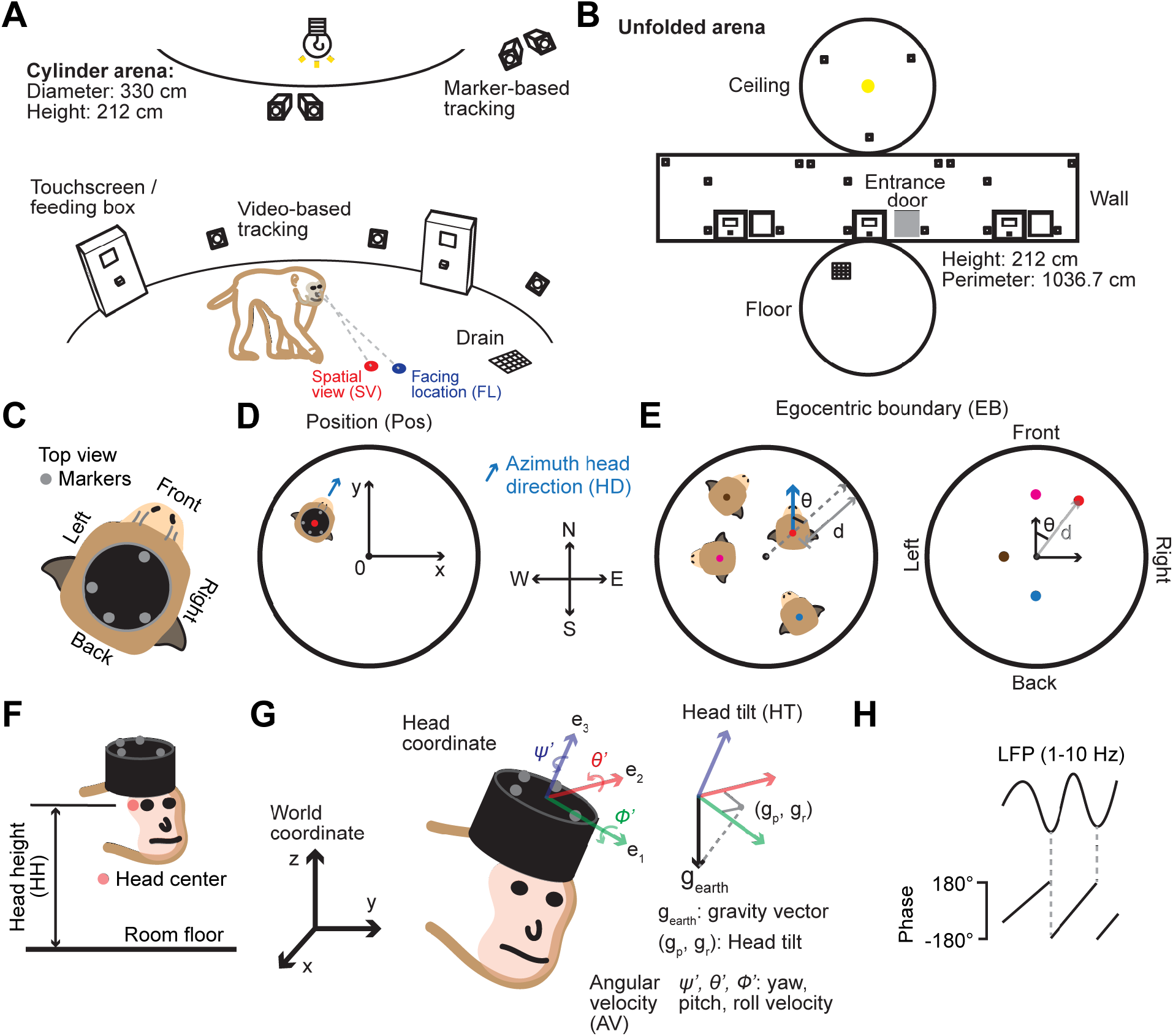
Experimental setup and behavioral variables, related to Figures 1-4. **(A)** Diagram of the circular arena (diameter: 330 cm; height: 212 cm), lit by a ceiling LED light, with touchscreen/feed boxes and drain as landmarks. Nine marker-based and nine video-based tracking cameras were placed on wall and ceiling. Monkeys entered/exited the arena through a sliding door. The ***Facing location (FL)*** variable was defined as the intersection between the heading vector (where the head points at) and the arena (blue blob on the floor). The ***Spatial view (SV)*** variable was defined as the intersection between the viewing (gaze) vector (where the left eye looks) and the arena (red blob). **(B)** Unfolded arena diagram used to define FL and SV tuning curves. **(C)** Top view of monkey head cap with 4 reflective markers for motion tracking. **(D) *Position (Pos)*** and ***Azimuth head direction (HD)*** were defined as the position/direction of the head in world coordinates (red dot/blue arrow). **(E) *Egocentric boundary (EB)*** was defined as the monkey’s distance (d) and HD angle (θ) relative to the closest point of the arena boundary, shown from allocentric (left) and egocentric (right) perspectives. Magenta/gray/blue dots: boundary in front (θ = 0°)/left (θ = 90°)/back (θ = 180°) of the animal. **(F) *Head height (HH)***, defined as the height of the head center (red dot) from the arena floor. **(G) *Head tilt (HT)***, defined as the projection of the gravity vector (g_earth_) onto the head axial plane, with components (g_p_, g_r_) corresponding to pitch and roll (thus, a 2D variable). ***Head angular velocity (AV)*** (yaw, pitch, and roll) was extracted from instantaneous head orientation. **(H) *Local-field potential phase***, extracted from band-pass filtered LFP (1-10 Hz), with troughs corresponding to ±180°.

**Figure S3.**
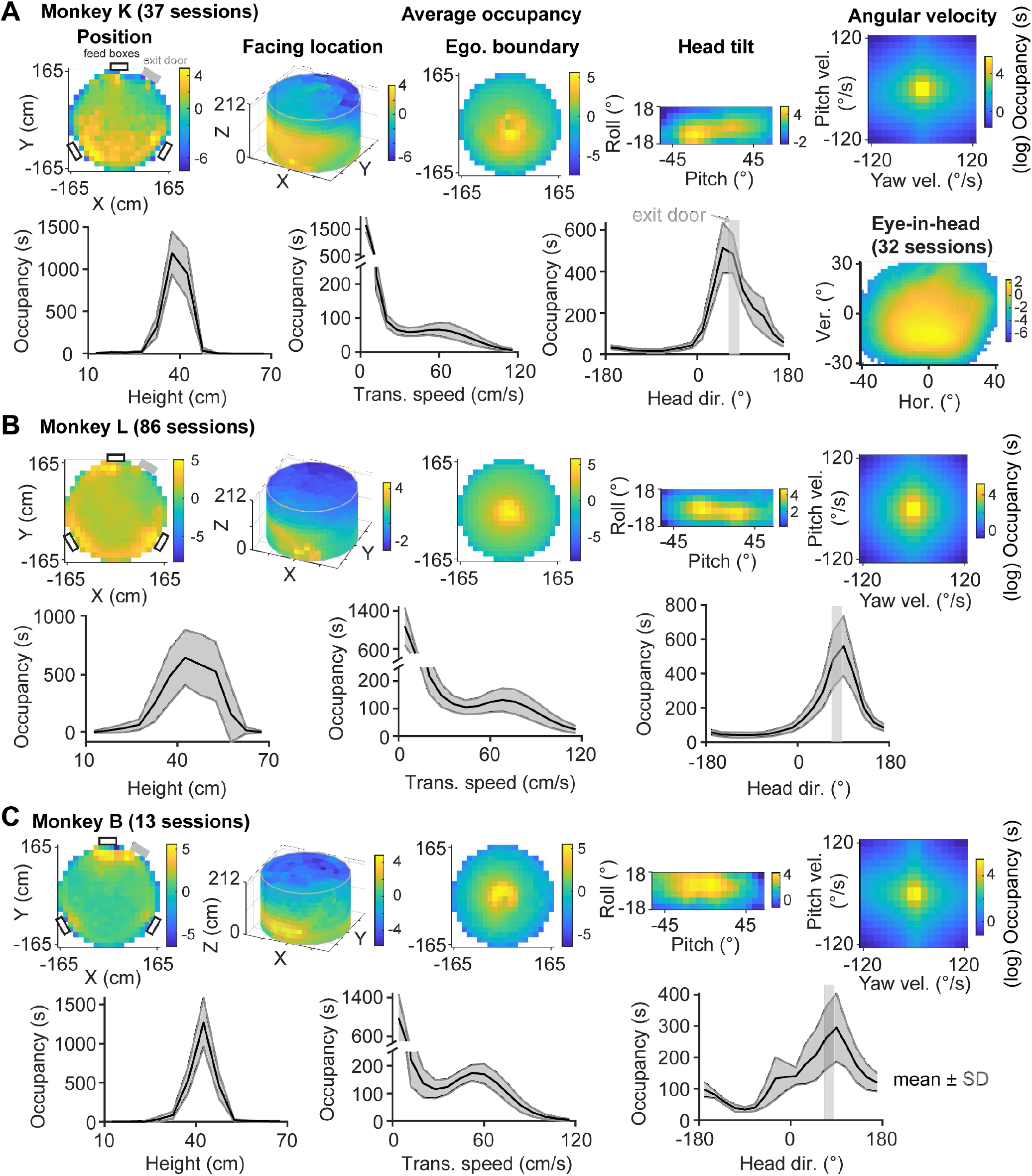
Occupancy maps of behavioral variables for each animal, related to Figures 2-4. **(A-C)** For 2D variables (Position, Head facing location, Egocentric boundary, Head tilt, Head angular velocity, and eye-in-head position), color indicates average time spent in each bin in natural logarithmic scale of seconds. Open rectangles: touchscreen/feed box locations. Gray rectangles: entrance/exit door location. For 1D variables (Head height, Speed, Azimuth Head Direction), shaded area corresponds to 1x standard deviation across sessions.

**Figure S4.**
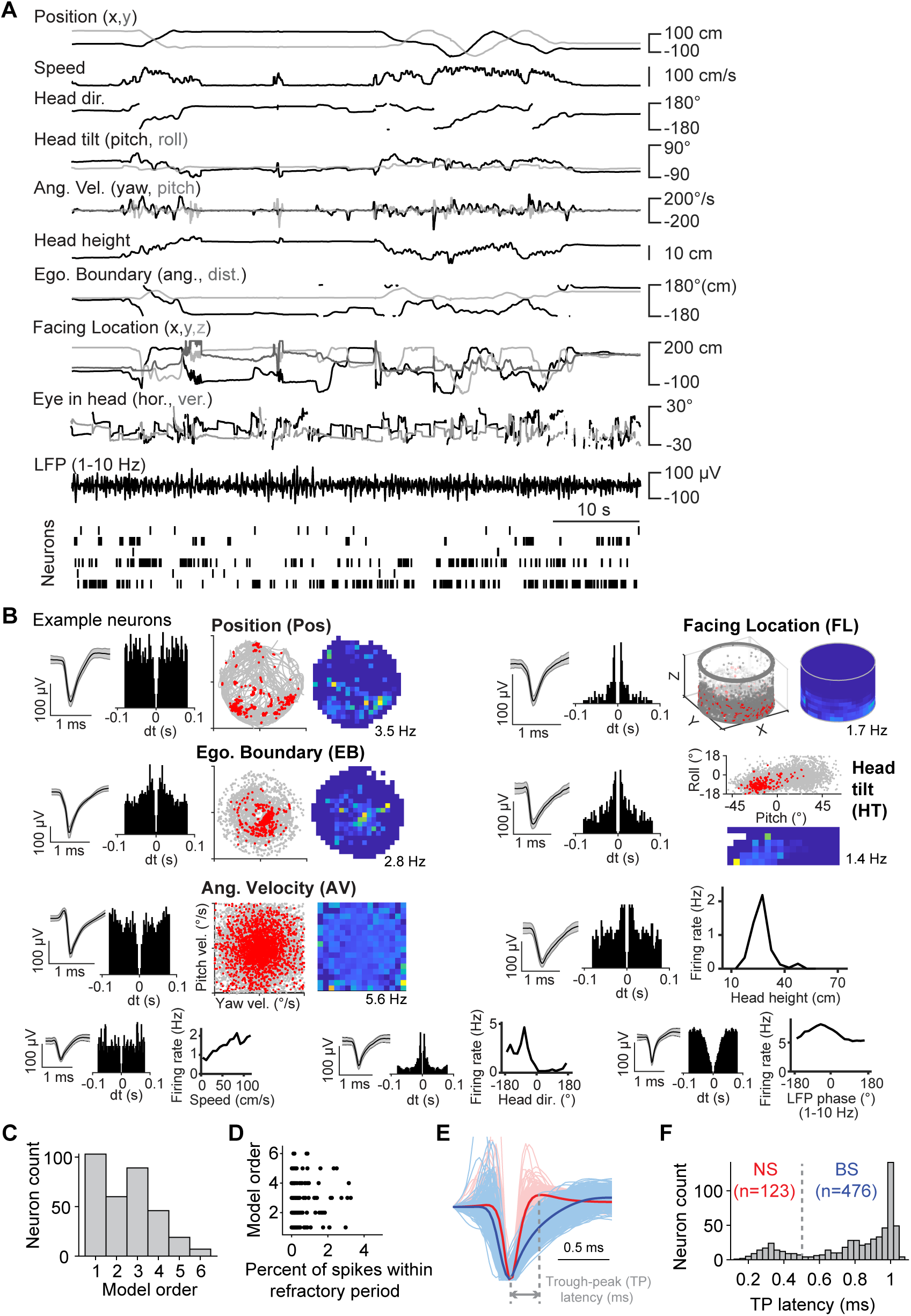
Neuronal tuning to diverse spatial variables, related to Figures 3 and 4. **(A)** A short segment of the raw traces of all behavioral variables and the corresponding spike raster. **(B)** Example neurons (from different sessions) with their raw tuning curves for select variables. From left to right: Average spike waveform (shaded area: 1x standard deviation), auto-correlogram, and tuning curves. For 2D tuning curves, down-sampled (for visualization purposes), two plots are shown: behavioral data (gray) with superimposed spikes (red dots), as well as raw firing colormaps. **(C)** Model order (the number of variables that are conjunctively encoded) distribution. **(D)** Model order vs. spiking sorting quality (refractory period violation). Neurons with 0 violation were not included. **(E)** Average spike waveforms for all neurons. Red, narrow-spiking (NS) neurons; blue, broad-spiking (BS) neurons. **(F)** Distribution of waveform trough-to-peak (TP) latency. NS, TP latency smaller than 0.5 ms; BS, TP latency larger than 0.5 ms.

**Figure S5.**
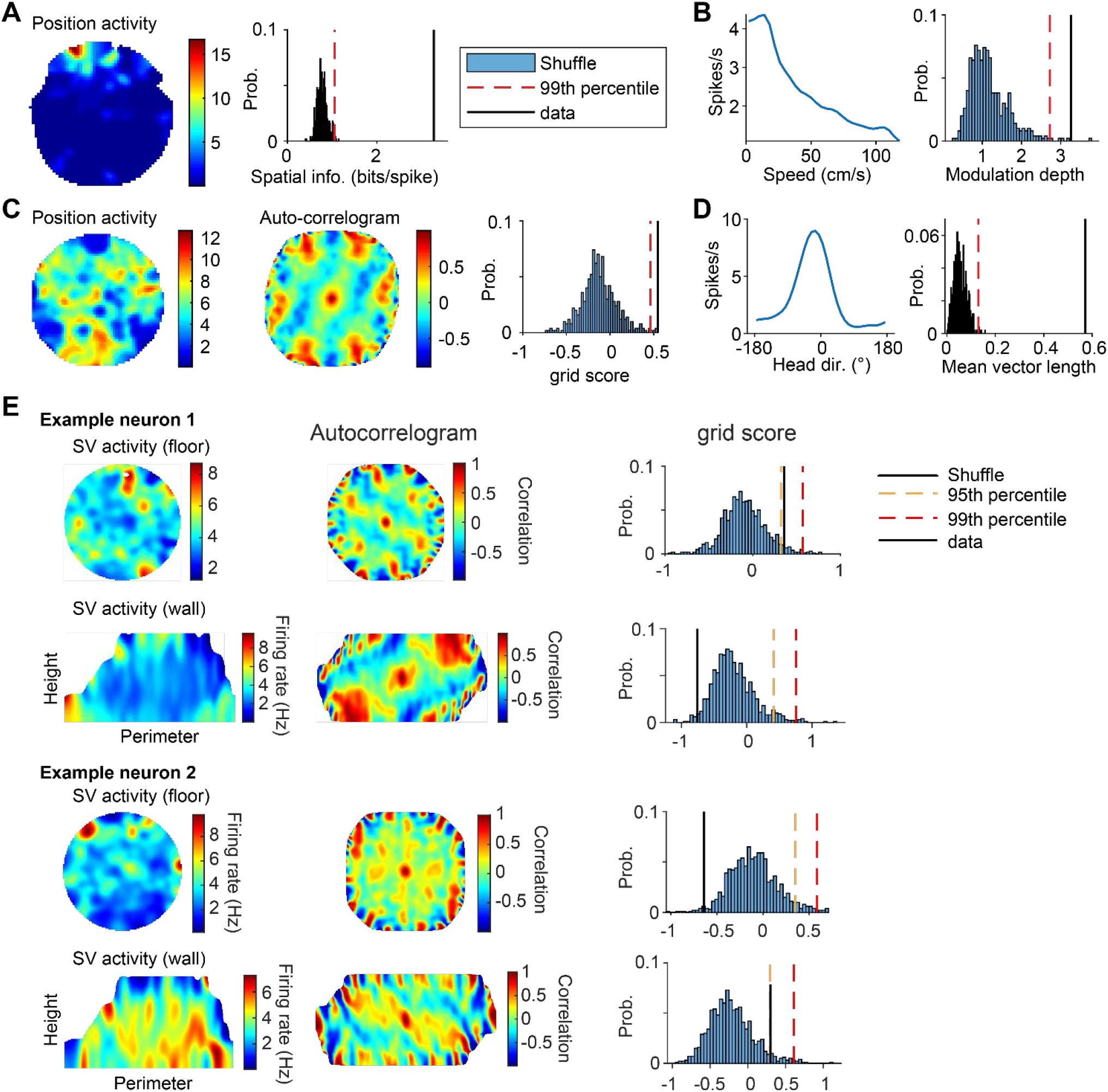
Spatial tuning analyses using traditional methods, related to Figures 3 and 7. **(A)** Left, position activity colormap for an example neuron. Right, spatial information for the actual data (black line) and shuffled distribution. Red dashed line indicates the 99^th^ percentile of the shuffled distribution. **(B)** Left, speed tuning curve for an example neuron. Right, modulation depth for the actual data and shuffled distribution. Modulation depth was calculated as the difference between the maximum and minimum firing rates in the speed tuning curve (Methods). **(C)** Left, position activity colormap for an example grid cell. Middle, auto-correlogram of the position activity map on the left. Right, grid score for the actual data and shuffled distribution. **(D)** Left, head direction tuning curve for an example neuron. Right, mean vector length for the actual data and shuffled distribution. **(E)** Grid tuning analysis for ‘spatial view’ (SV) variable. Two example neurons are shown. Left, SV activity on the floor and wall. Middle, auto-correlogram of the corresponding SV activity map on the left. Right, grid score for the actual data and shuffled distribution; 95^th^ and 99^th^ percentiles are shown. Note no neurons passed the 99^th^ percentile criterion.

**Figure S6.**
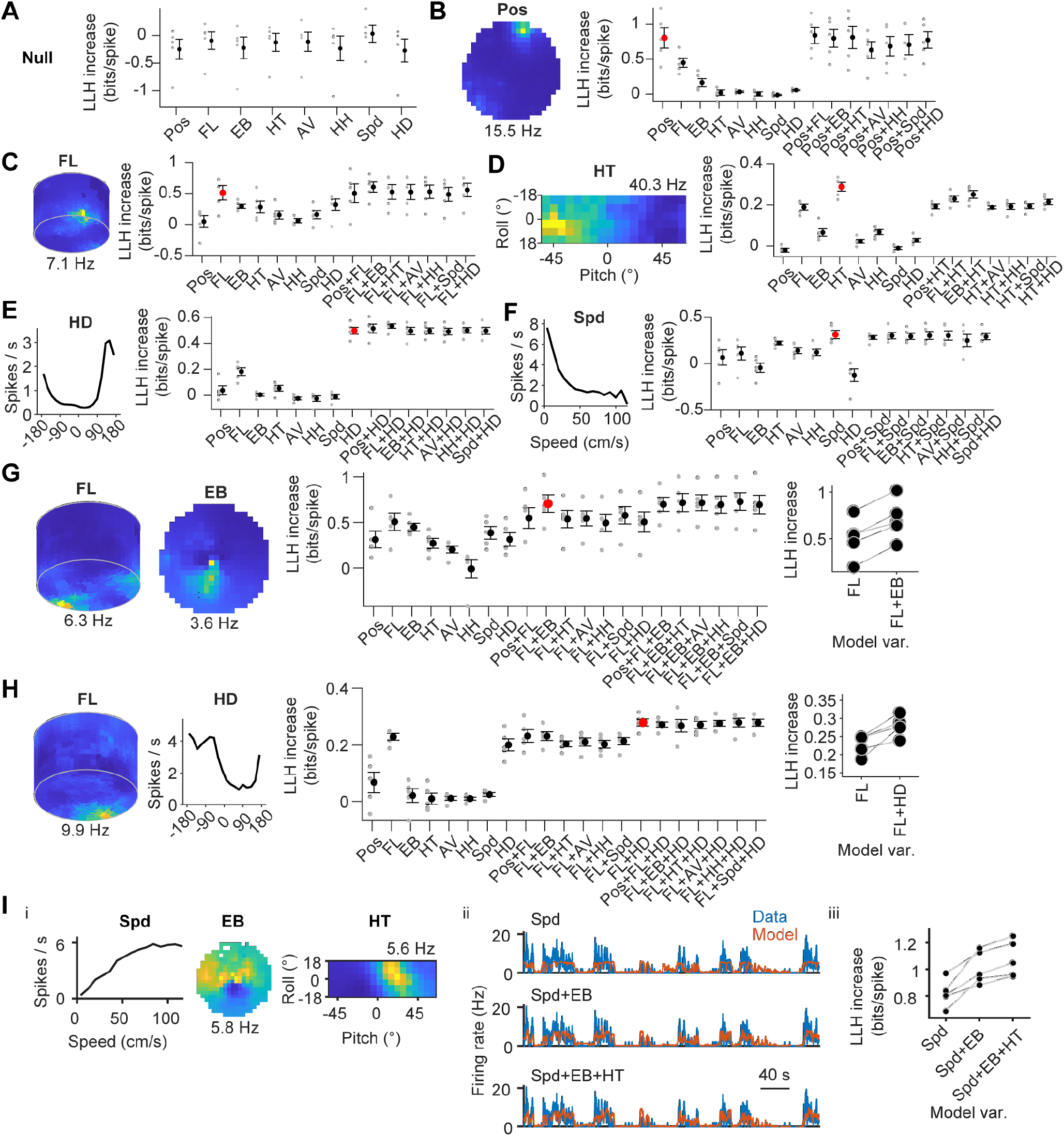
Example model-based tuning curves and model selection, related to Figure 4. A forward search approach was used to identify the best model. Black dots: increase of the mean log-likelihood (LLH) (from the null, mean firing rate model). A LLH increase not significantly larger than 0 indicates that the current model does not perform better than the null model. Red dot: best model selected. **(A)** An example neuron not tuned to any variable. All LLH values of the single-variable (1^st^ order) models are not significantly larger than 0. **(B-F)** Example neurons tuned to a single variable, with model-based tuning curves shown on the left. Examples tuned to position (Pos), facing location (FL), head tilt (HT), azimuth head direction (HD), and speed. **(G) and (H)** Example neurons tuned to combination of 2 variables, with model-based tuning curves shown on the left. Examples tuned to FL+EB and FL+HD. **(I)** An example neuron whose activity was best fitted by a 3^rd^-order model: a combination of speed, egocentric boundary, and head tilt. (i) Model-based tuning curves. Peak firing rates of the colormaps are indicated (lowest was 0). (ii) Actual (blue) and fitted (orange) firing rate shown for a short segment for the 1^st^, 2^nd^ and 3^rd^-order models. (iii) LLH increase as a function of model variables. Each line corresponds to one-fold in the cross-validation process.

**Figure S7.**
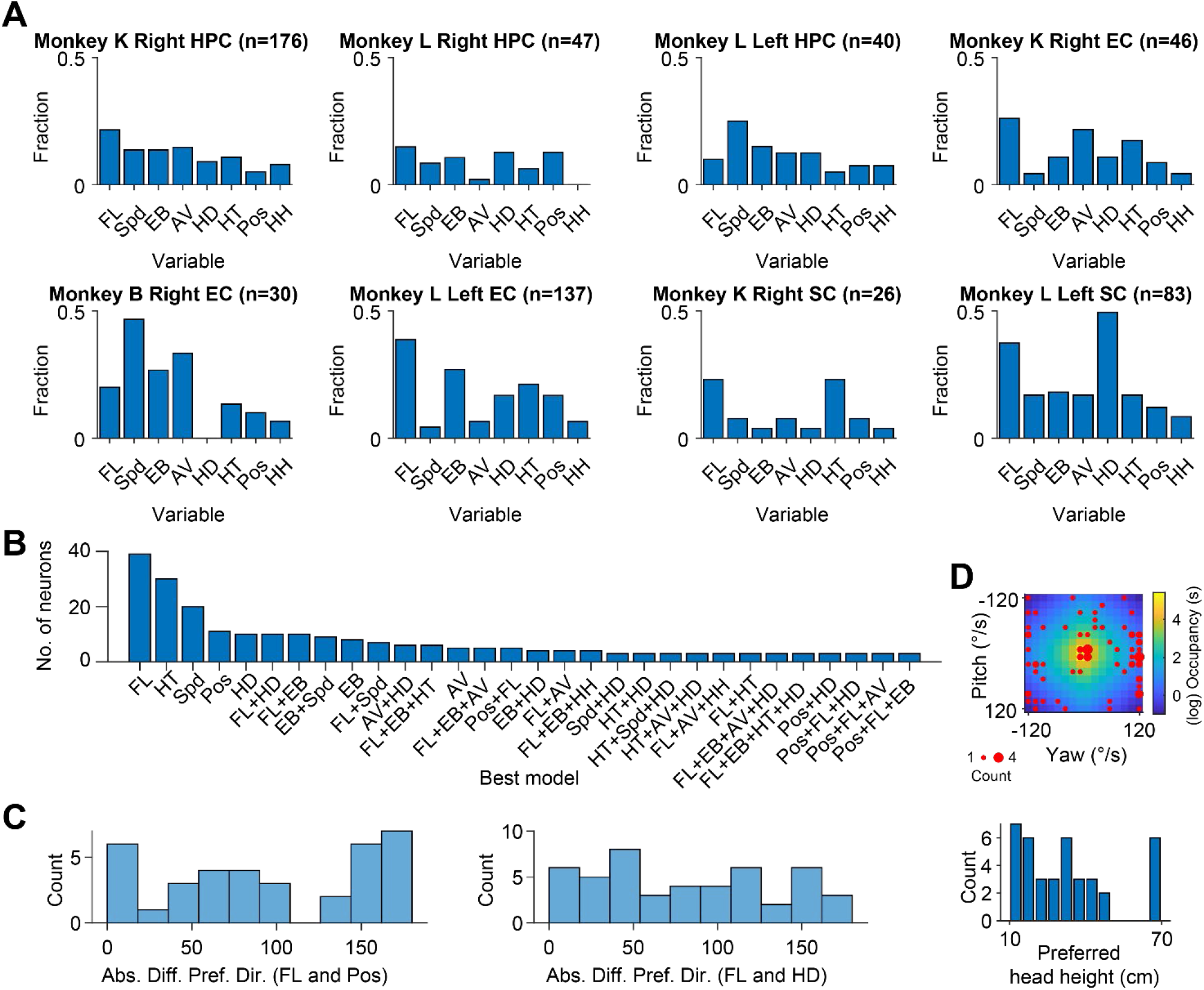
Spatial tuning across monkeys and regions, related to Figures 4 and 5. **(A)** Breakdown of the fraction of encoding neurons in individual regions, hemispheres, and monkeys. **(B)** Neuron count captured by the top 30 best models across regions. The best model variables are indicated as the x tick labels. **(C)** Left: distribution of the absolute difference between the preferred direction (in room coordinates) for variables FL and Pos (using neurons tuned to both FL and Pos). Right: distribution of the absolute difference between the preferred direction for variables FL and HD (using neurons tuned to both FL and HD). **(D)** Top, preferred firing fields (red dots) for all neurons encoding AV superimposed on the average occupancy colormap across all monkeys. Dot size corresponds to neuron count. Bottom, distribution of preferred head height for all tuned neurons.

